# Evaluating the quorum quenching potential of bacteria associated to *Aurelia aurita* and *Mnemiopsis leidyi*

**DOI:** 10.1101/602268

**Authors:** Daniela Prasse, Nancy Weiland-Bräuer, Cornelia Jaspers, Thorsten B.H. Reusch, Ruth A. Schmitz

## Abstract

The associated microbiota of marine invertebrates plays an important role to the host in relation to fitness, health and homeostasis of the metaorganism. As one key chemically-mediated interaction, Quorum sensing (QS) and interference with QS among colonizing bacteria ultimately affects the establishment and dynamics of the microbial community on the host. Aiming to address interspecies competition of cultivable microbes associated to merging model species of the basal animal phyla Cnidaria (*Aurelia aurita*) and Ctenophora (*Mnemiopsis leidyi*) as well as to evaluate their potential to shape the associated community by interfering with QS, we performed a classical isolation approach. Overall, 84 bacteria were isolated from *A. aurita* medusae and polyps, 64 bacteria from *M. leidyi*, and 83 bacteria from the ambient seawater, followed by taxonomically classification by full length 16S rRNA gene analysis. The results show that the moon jellyfish *A. aurita* and the comb jelly *M. leidyi* harbor a cultivable core microbiota consisting of typical marine and ubiquitously found bacteria (e.g. *Chryseobacter, Microbacterium, Micrococcus, Olleya, Phaeobacter, Pseudoalteromonas, Pseudomonas, Rhodococcus, Shewanella, Staphylococcus*, and *Vibrio*) which can also be found in the ambient seawater. However, several bacteria were restricted to one host (e.g. for *A. aurita: Bacillus, Glaciecola, Ruegeria, Luteococcus;* for *M. leidyi: Acinetobacter, Aeromonas, Colwellia, Exiguobacterium, Marinomonas, Pseudoclavibacter, Psychrobacter, Sagittula, Thalassomonas*) suggesting host-specific microbial community patterns. Evaluating QQ activities, out of 231 isolates, 121 showed QS-interfering activity. They mainly interfered with the acyl homoserine lactone (AHL) based communication, whereas 21 showed simultaneous quorum quenching activities against AHL and autoinducer-2. Overall, this study provides insights into the cultivable part of the microbiota associated to two environmentally important marine non-model organisms and discloses their potential in synthesizing QS interfering compounds, potentially important in shaping a healthy and resilient microbiota.

## Introduction

The marine environment covers more than 70 % of the world’s surface and harbors approximately 3.6 × 10^28^ microorganisms (1–4). Marine microbial communities are highly diverse and have evolved under a variety of ecological conditions and selection pressures (5). Those microbes are also known to form beneficial symbiotic relationships with various marine multicellular hosts, e.g. sponges, corals, squids, and jellies, and are assumed to produce unique biologically active compounds important for the homoeostasis of the host, which are often pharmacologically valuable compounds (6, 7). Recently, the potential functional role of associated microbiota became an important research focus in the fields of zoology, botany, ecology and medicine, since each multicellular organism can be regarded as an entity with its associated microbes as a so-called ‘metaorganism’, where the microbes most likely have crucial functions for the fitness and health of the host (6, 7). The role of bacteria during the development of various organisms, such as humans, as well as their importance for the host’s resilience in the control of pathogens and prevention of inflammatory diseases has been intensively studied in recent years reviewed in (8). These studies also showed that the interactions within a metaorganism are considerably complex. In order to understand this complexity, research studies dealing with the impact of the microbiota on a host have to access already well-studied model organisms, such as the *Drosophila* and mouse model. To understand the long-term evolutionary origin of the metaorganism, however, novel model organisms are urgently needed. Here, evolutionarily ancient organisms, which are located at the base of the metazoan tree, will provide important insight. They are often widely distributed in the marine environment because of their enormous adaptability, they have a simple morphology with only two tissue layers as surfaces for microbial colonization, and some of them have complex life cycles with alternating life stages (6). Two marine basal metazoans, which combine these characteristics, are the moon jellyfish *Aurelia aurita* and the comb jelly *Mnemiopsis leidyi* (Fig. 1A). *A. aurita* is one of the most widely distributed Scyphozoans (Cnidaria) featuring a complex life cycle. In its di-phasic life cycle, *A. aurita* alternates between a free-living pelagic medusa and a sessile polyp. In its different successive stages of life cycle - planula larva, sessile polyp, pelagic ephyra and medusa - *A. aurita* is adapted to different marine environmental factors, e.g. salinity, temperature, and food spectrum (9). In a recent study, we disclosed that those different life stage and moreover different compartments of a medusa as well as different sub-populations harbor distinct, specific microbiota (10). On the other hand, the comb jelly *M. leidyi* (Ctenophora) is one of the most successful invasive marine species worldwide (11). *M. leidyi* features an unusual mode of reproduction so-called dissogony (12), where small larval stages are already sexually reproducing. Adults are reproductively active again and are simultaneous hermaphrodites (12). Although those jellies possess only two tissue layers for bacterial colonization, interactions at those interfaces can be manifold and take place between the host and the colonizing bacteria and among the bacteria. One essential interaction between bacteria regulating population behavior is the bacterial cell-cell communication, so-called quorum sensing (QS). Most bacteria are capable to synthesize and secrete small signal molecules (autoinducers), which are detected to respond to changing cell population densities (13, 14). Examples for coordinated population behavior based on QS are bioluminescence, virulence, pathogenicity, biofilm formation and colonization (13, 14). Consequently, these population behaviors can be affected by molecules interfering with QS (quorum quenching (QQ) molecules). Communication interfering compounds have been described to be produced and secreted by competing bacteria, but host-derived interference has also been reported (15–17). QS and the respective interference might be important to maintain the healthy stability of the microbiota and the metaorganism, and prevent colonization by pathogens (15–17).

**Fig. 1:**
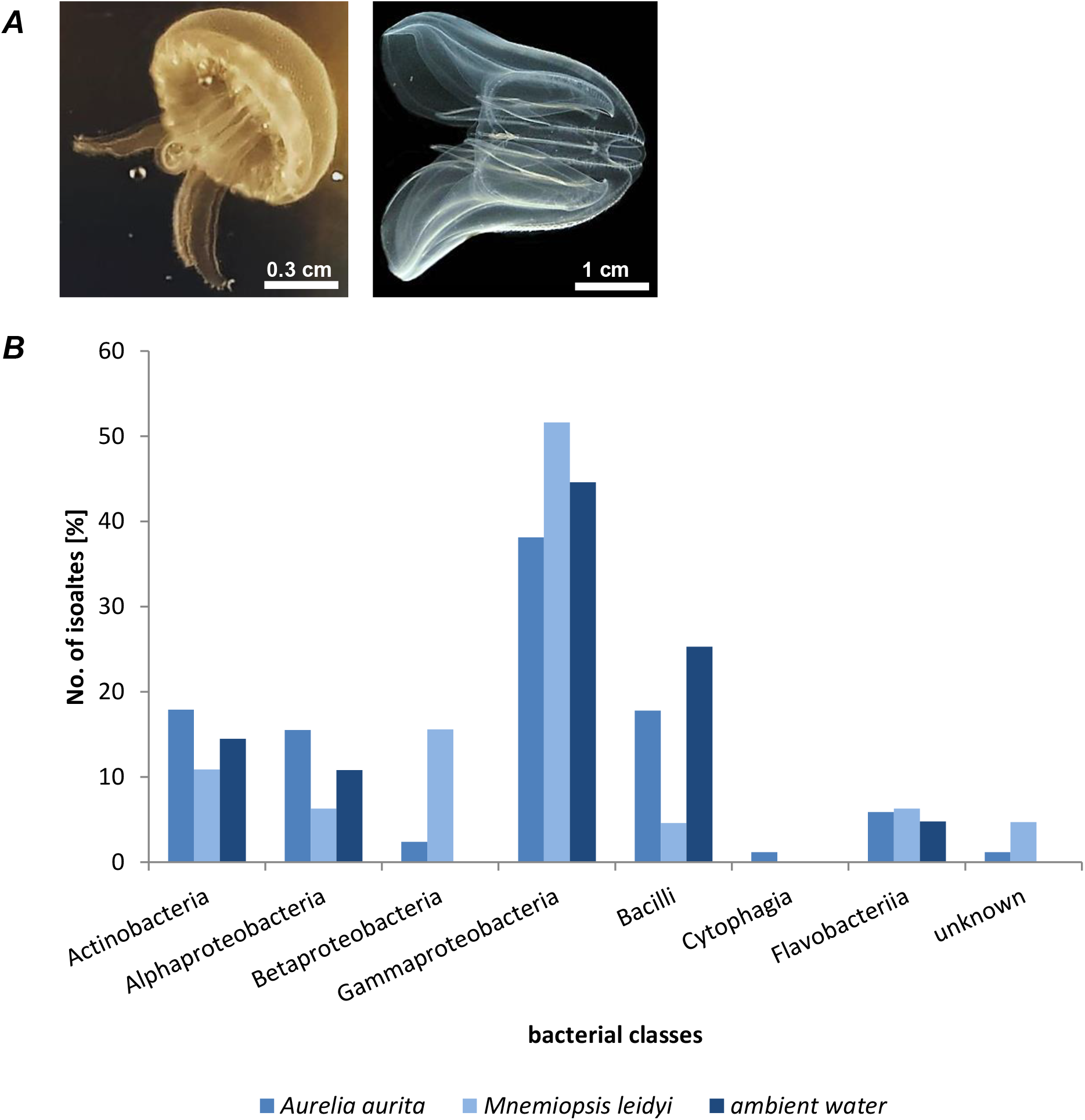
Taxonomic classification of isolates. (***A***) *Aurelia aurita* (left panel) and *Mnemiopsis leidyi* (right panel) (***B***) Bar plot represents classification of bacterial isolates to class level in percentage.

Current molecular high-throughput techniques, like deep sequencing, are preferred methods to analyze such complex microbial communities and interactions as those occurring on the above mentioned jellies (18). They enable the identification of microbes, normally missed by cultivation-based approaches and ultimately allow for a broader picture of the entire environmental network (18). However, these metagenomic-based approaches lack the possibility for experimental manipulation studies. Although the cultivation of certain bacteria is still challenging, several recent studies showed that the cultivation of host-associated bacteria is urgently needed to better study and understand their function in a metaorganism (19). Thus, this study aimed to isolate bacterial colonizers of *A. aurita* and *M. leidyi* to disclose potentially specific microbial community patterns of the jellies compared to the ambient seawater. Another intention was to elucidate the frequency of QQ active bacteria within those isolates, which on the one hand might be important for the maintenance of a healthy microbiota by defending competitors or pathogens, and on the other hand might be a rich source for biotechnologically relevant compounds.

## Material and methods

### Sampling of *A. aurita* and *M. leidyi*

Individual *A. aurita* (Linnaeus) and *M. leidyi* medusae (with mean size umbrella diameter of 22 cm and 4 cm, respectively) were sampled from one location in the Kiel Bight, Baltic Sea (54°32.8’N, 10°14.7’E) in June 2015 and May 2017 using a dip net. The animals were immediately transported to the laboratory and washed thoroughly with autoclaved artificial seawater to remove non-associated microbes. Additionally, medusae were kept in artificial seawater (Tropical Marine Salts, 18 practical salinity units (PSU)) for ten days before thoroughly washing with autoclaved artificial seawater to remove non-associated microbes and considered as husbandry medusae. Moreover, 1L ambient water (Baltic Sea 54°32.8’N, 10°14.7’E; artificial seawater 18 and 30 PSU) was filtrated (0.22 μm pores, Millipore, 45 mm diameter) to enrich bacterial cells for further isolation. *A. aurita* polyps of sub-populations Baltic Sea, North Sea and North Atlantic (Roscoff) were kept in artificial seawater (Tropical Marine Salts, 18 PSU (Baltic Sea) and 30 PSU (North Sea, North Atlantic)) in 2 L plastic tanks. Prior to bacterial enrichment and isolation, polyps were not fed with freshly hatched *Artemia salina* for at least three days.

### Bacterial enrichment and isolation

Bacterial enrichment and isolation was performed with cotton swabs from the surface of the umbrella of native and kept *A. aurita* as well as *M. leidyi* medusae, whole homogenized *A. aurita* polyps of different sub-populations, and filters derived from ambient seawater filtration. In order to ensure high diversity during the isolation procedure, swabs, 100 μL of homogenized animal tissues, and ¼ of a filter were streaked/plated onto three different solid media (Marine Bouillon (20), R2A (21), Plate Count medium (22)) and incubated at 4 °C, 20 °C ad 30 °C. Obtained single colonies were picked and purified by streaking at least three times on the respective agar plates. Morphologically different colonies from the different sample types were grown in the respective liquid medium at the respective incubation temperature. Pure cultures were stored as glycerol stocks (10 %) at – 80 °C and were subsequently taxonomically classified by 16S rRNA analysis.

### Nucleic acid isolation

High-molecular weight genomic DNA of bacteria was isolated from 5 mL overnight cultures using the Wizard genomic DNA isolation kit (Promega) according to the manufacturer’s instructions. Fosmid DNA was isolated from 5 mL overnight cultures using the Presto Mini Plasmid kit (Geneaid) according to the manufacturer.

### 16S rRNA gene analysis

16S rRNA genes were PCR amplified from 50 ng isolated genomic DNA using the bacterium-specific primer 27F (5′-AGAGTTTGATCCTGGCTCAG-3′) and the universal primer 1492R (5′-GGTTACCTTGTTACGACTT-3′) (23) resulting in a 1.5-kb bacterial PCR fragment. The fragment sequences were determined by the sequencing facility at the Institute of Clinical Molecular Biology, University of Kiel, Kiel, Germany (IKMB).

### Construction of a genomic large-insert library

A genomic library of *Pseudoalteromonas issachenkonii* (isolate no. 91) was constructed using the Copy Control Fosmid Library Production Kit and vector pCC1FOS (Epicentre) with the modifications described by Weiland et al. (24).

### Identification and molecular characterization of quorum sensing interfering clones

Fosmid clones were grown separately in 200 μL LB medium in microtiter plates. Preparation of cell-free culture supernatants and cell extracts of pools of 96 fosmid clones and individual clones of QQ positive 96er pools was performed as described in (25). QQ assays on plates using strains AI1-QQ.1 and AI2-QQ.1 were performed with cell-free supernatants and cell extracts of fosmid clones and purified proteins as previously described in (25). In order to identify the respective ORFs of the fosmids conferring QQ activity, subcloning was used as previously described in (25, 26). Putative QQ-ORFs were PCR-cloned into pMAL-c2X N-terminally fusing the QQ-ORFs to the maltose binding protein (MPB) using ORF-specific primers adding restriction recognition sites flanking the ORFs (see Tab. S1); overexpressed and proteins purified as recently described in (26).

## Results and Discussion

### Isolation of jelly-associated bacteria conforming host specific microbiota

The overall goal of this study was to enrich and isolate bacterial colonizers of the moon jellyfish *A. aurita* and the comb jelly *M. leidyi* to unravel potential host-specific patterns in their microbial composition based on a cultivation-dependent approach. These isolates will be later on used in controlled laboratory experiments to study their function in the metaorganism in more detail (e.g. by recolonization of germ-free hosts with manipulated microbial communities). Moreover, we aimed to gain insights into the potential of those isolates in synthesizing interesting novel biomolecules, particularly QQ active compounds. On the one hand, those might be important for establishing and maintaining a healthy microbiota of the metaorganism, and on the other hand might act as biotechnologically relevant compounds. Thus, a classical isolation approach was performed with three different solid media at three different incubation temperatures to ensure high diversity in enriched bacteria. Plate Count Agar was used for enrichment of the viable bacterial fraction of a sample without any selection, whereas R2A was used for enrichment of heterotrophic, aquatic bacteria, which tend to be slow-growing species and might be suppressed by faster-growing species on a richer culture medium, and Marine Bouillon was used to select for marine bacteria preferring high-salt conditions (see Materials and methods).

In total, we isolated 84 bacterial strains from *A. aurita* (Tabs. 1+ Table S2). In more detail, 17 bacteria were isolated from the umbrella surface of native Baltic Sea medusae and 8 bacteria from cultured medusa. From the benthic polyps cultured in the laboratory, 22 bacteria were isolated from the Baltic Sea sub-population, 19 from the North Sea sub-population, and 18 from the North Atlantic sub-population (Tabs. 1+ Table S2). The isolation procedure from *M. leidyi* resulted in the identification of 64 bacteria (Tabs. 1+ Table S2), 36 bacteria were enriched from native Baltic Sea medusae and 28 from cultured ones. Moreover, 83 bacteria were isolated from the ambient seawater (Tabs. 1+ Table S2). Full length 16S rRNA gene analyses revealed the identification of eight different bacterial classes (Fig. 1), representing 28 families and 51 genera. Overall, distinct differences in the microbial composition were observed between ambient seawater and the two jellies (summarized in Fig. 1, Tabs. 1+ Table S2). In the cultivation approach, representatives of the classes Betaproteobacteria, Cytophagia and unclassified bacteria were shown to be present on jellies, but were not isolated from the surrounding seawater samples (Fig. 1), whereas several bacteria were exclusively isolated from the ambient seawater and not from the jellies and assigned to the genera *Celeribacter, Corynebacterium, Fictibacillus, Halomonas, Leisingera, Maribacter, Marinobacter, Salinibacterium* and *Serratia* (Tabs. 1+ Table S2, Fig. 2). All of those bacteria are typically found in the marine environment, some of them also associated with marine eukaryotes (10, 16, 27, 28). These cultivation-dependent results are in line with several publications demonstrating that the microbiota associated to animals is significantly different to the bacterial communities in the ambient seawater (10, 16, 27, 28). Specific selection mechanisms, both on bacterial as well as on host site, have to occur to establish and maintain such a specific host microbiota (29, 30). Notable are the frequencies with which the ubiquitous and high abundant occurring bacteria of the genera *Pseudoalteromonas* (violet) and *Pseudomonas* (orange) were isolated from all samples (Fig. 2). These occur in open waters but are also able to colonize animal tissues (31, 32). Both genera are well-known marine inhabitants found in symbiotic associations with sponges, corals and algae, and additionally are known for their versatile biotechnological potential with respect to the production of antimicrobials and enzymes of industrial interest (33–35).

**Fig. 2:**
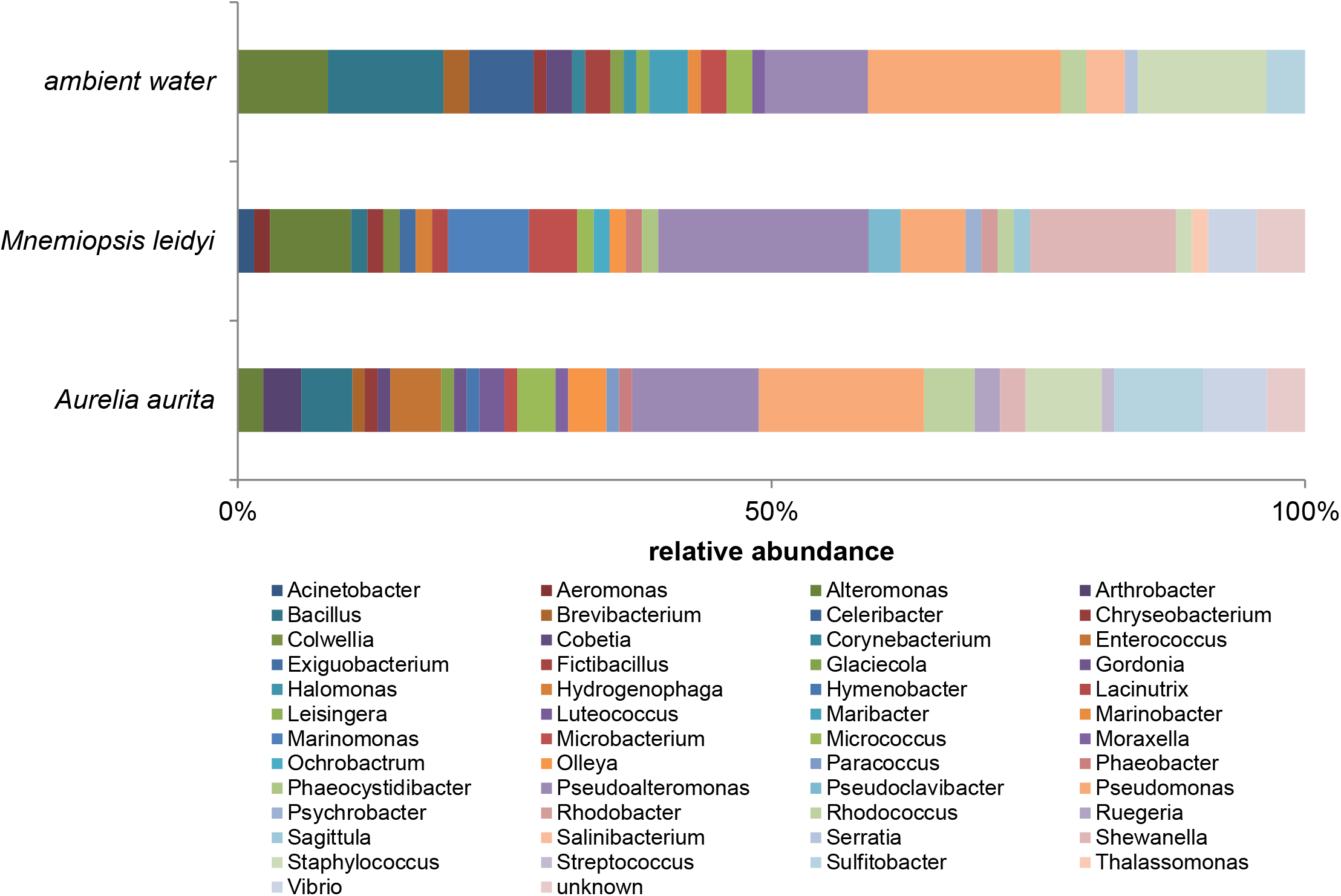
Taxonomic classification of isolates. Bar plot represents relative abundance of isolates in the different sample types based on genus level.

**Tab. 1:**
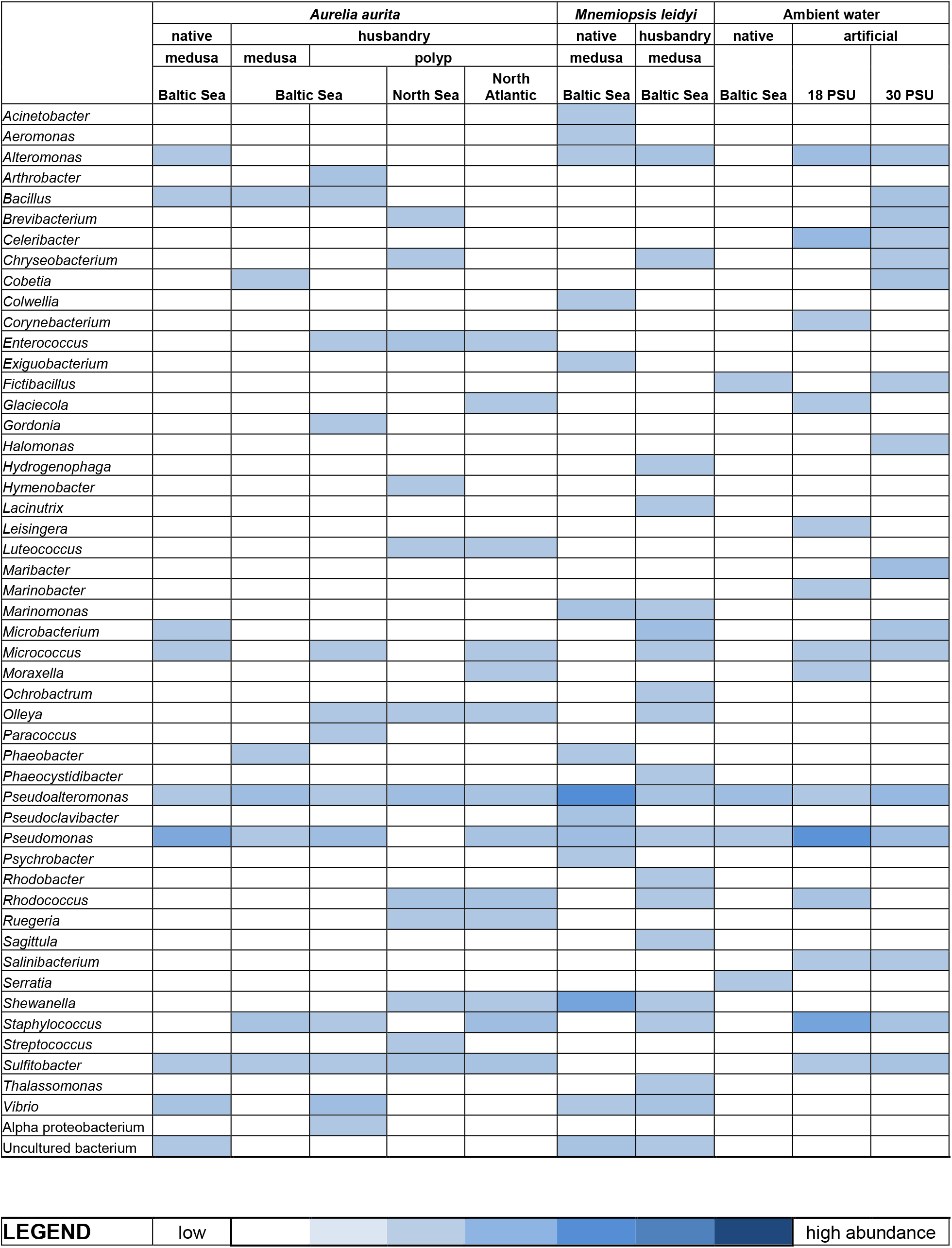
Presence of isolates in different sample types. Presence and abundances of isolates in different sample types is visualized as heat map (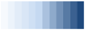 low → high abundance) on genus level.

Phylogenetic classification revealed that *A. aurita* is colonized by cultivable representatives of genera *Arthrobacter, Bacillus, Brevibacterium, Cobetia, Enterococcus, Glaciecola, Gordonia, Hymenobacter, Lutococcus, Paracoccus, Phaeobacter, Ruegeria, Streptococcus* and *Sulfitobacter* (Tab. 1+S2, Fig. 2). In more detail, *Bacillus* was exclusively isolated from Baltic Sea specimens, *Enterococcus* from benthic polyps, and *Glaciecola, Ruegeria* and *Luteococcus* from high-salt kept (30 PSU) polyps (Tab. 1). Regarding an important function of those bacteria for the host can only be speculated, but the Tropodithietic acid-producing genus *Ruegeria* of the Roseobacter clade is a globally distributed marine bacterial species found primarily in the upper open ocean and has primarily been isolated from marine aquaculture, where *Ruegeria* strains have probiotic potential (36). Additionally, *Luteococcus* has been isolated from the marine environment and was often found in the intestinal tracts of animals (37). A highly specific colonization of *A. aurita* was already shown in a 16S rRNA deep sequencing study demonstrating jellyfish-specific microbial patterns, which were even different in different compartments of a medusa, between different sub-populations, and among different life stages (10). In the present study, our cultivation-dependent data also indicate such host-specific community patterns. All bacteria isolated from *A. aurita* have been found in the deposited 16S data sets (10) suggesting that the isolated bacteria indeed reflect the cultivable proportion of the moon jellyfish microbiota. We were able to cultivate a high proportion of representatives (14 out of 24 genera) identified by 16S amplicon sequencing.

Such a specificity in microbial communities can also be suggested for *M. leidyi*, where likewise, a diverse set of associated colonizers was isolated, which were assigned to the genera *Acinetobacter, Aeromonas, Alteromonas, Colwellia, Exiguobacterium, Hydrogenophaga, Lacinutrix, Marinomonas, Moraxella, Ochrobactrum, Phaeocystidibacter, Pseudoclavibacter, Pychrobacter, Rhodobacter, Sagitulla* and *Thalassomonas* (Tab. 1+ Table S2, Fig. 2). Bacteria belonging to the genera *Chryseobacter, Microbacterium, Micrococcus, Olleya, Phaeobacter, Pseudoalteromonas, Pseudomonas, Rhodococcus, Shewanella, Staphylococcus* and *Vibrio* were isolated from both jellies and most of them also from the ambient seawater (Tab. 1, Fig. 2). These bacteria most likely are ubiquitous marine bacteria, whose abundances in the open waters might differ from their abundances on the animals. As both jellies share the same environment and use the same food sources, it is not astonishing that they also share a core marine microbiota. Nonetheless, several bacteria were exclusively isolated from *M. leidyi*, such as *Acinetobacter, Aeromonas, Colwellia, Exiguobacterium, Marinomonas, Pseudoclavibacter, Psychrobacter, Sagittula*, and *Thalassomonas* (38). Remarkably, all those representatives were not isolated from the ambient seawater suggesting that those bacteria are host-specific and potentially involved in the invasion process of the comb jelly (39–41). Several bacteria were previously described in connection with *M. leidyi*. For instance, *Aeromonas* was already isolated from *M. leidyi* of the North Sea population (41), *Marinomonas* was even described as the dominant phylotype of the comb jelly in a 16S amplicon-based study (39), and *Colwellia* was isolated from several marine animal and plant tissues (27). Further, *Colwellia* has already been identified on various brown algae and the red alga *Delisea pulchra* using shotgun sequencing, where it was present on diseased thalli and absent from healthy thalli of this red alga (42). The hypothesis that those bacteria indeed play crucial roles for the comb jelly, in particular during the invasion process, has to be proven with a cultivation-independent approach.

Taken together, our cultivation-dependent approach revealed that the moon jellyfish *A. aurita* and the comb jelly *M. leidyi* harbor a core microbiota consisting of typically marine and ubiquitous bacteria, which can be also found in the ambient seawater, but in different abundances. Those bacteria were often found in microbiotas of other marine organisms, such as brown algae or fish gut (16, 27, 28). In addition, several bacteria were exclusively isolated from one of the jellies suggesting that the animals have their individual host-specific microbial communities, even if they share the same environment. The metaorganism, as entity of the host and its microbiota, has to control and modulate the microbial colonization of the host tissues to establish and maintain the specific microbiota, which most likely contributes to the overall fitness and health of the metaorganism.

### Screening for novel quorum quenching molecules

One important mechanism to control bacterial colonization is the interference with bacterial cell-cell communication, so called quorum sensing (QS) (13). QS is the fundamental system of bacteria regulating cell-density dependent behaviors, like virulence, pathogenicity, biofilm formation and colonization. QS is based on the synthesis and detection of small signal molecules called autoinducers (43). In Gram-negative bacteria, N-acetyl-homoserine-lactones (AHL) are synthetized by LuxI and bound by LuxR after reaching a given threshold (“quorum”). LuxR undergoes a change in conformation and acts as a transcription activator for target genes (14, 44). Gram-positive bacteria use short oligopeptides for QS-regulation (45). Besides this intra-specific communication, AI-2 – a furanone synthesized by LuxS - acts as autoinducer in the universal communication among different species (46–48). QS can be interfered by so-called quorum quenching (QQ) molecules, which for instance inactivate autoinducer-synthetases, degrade or modify the autoinducers or interfere with the autoinducer receptors through signal analogues. QQ molecules are synthesized by various bacteria. Ultimately, QS-regulated coordinated behaviors like colonization can be affected by the QQ molecules and might have important consequences in shaping the behavior and structure of polymicrobial communities. Many QS-regulated products are shared "public goods” that can be used by any member of the bacterial community (49), e.g. secreted proteases (50, 51). The production of sufficient "public goods” can only be enabled when the presence of cheating bacteria is hold at bay by interfering them through QQ (52–55). Moreover, many bacterial species use QS to control the production of toxins: for example, bacteriocins in *Streptococcus* species (56, 57) and type VI secretion effectors in *B. thailandensis* (58). Here, QQ prevents the increase of potential pathogens and the detrimental altering of community dynamics.

Thus, to evaluate the impact of QQ all isolated bacteria were screened for QS-interfering activities using the established reporter strains AI1-QQ.1 and AI2-QQ.1 (25). Screening cell-free cell extracts and culture supernatants demonstrated that 121 out of the 231 isolated bacteria showed AHL-interfering activities (52 %), of which 21 (9 %) showed simultaneous interference with AHL and AI-2 (Tab. S2, Fig. 3A). It is notable that over 50 % of the isolates evaluated showed QQ activities demonstrating a high frequency of QQ-active bacteria in microbial consortia associated with marine eukaryotes. One bacterium isolated from a polyp of sub-population Baltic Sea (no. 91, *Pseudoalteromonas issachenkonii*) exhibiting high QQ activities against AHL and AI-2 was selected for further molecular characterization. A genomic fosmid library was constructed to identify the respective ORF(s) conferring QQ activity. Cell extracts and culture supernatants of 480 individual fosmid clones were screened for QQ activity using the reporter strains AI1-QQ.1 and AI2-QQ.1 resulting in the identification of one fosmid (clone 5/E6) conferring the QQ activities simultaneous against AHL and AI-2. The identification of the respective ORF(s) mediating the QQ activities of the fosmid clone was achieved by subcloning followed by sequence analysis. In total, 3 putative QQ-ORFs were identified, which were PCR cloned into the pMAL-c2X expression vector resulting in QQ proteins N-terminally fused to the Maltose Binding Protein (MBP) (see Supplemental data set 1). Overexpressed MBP-QQ fusion proteins were purified by affinity chromatography and evaluated regarding their quenching activity using the reporter strains AI1-QQ.1 and AI2-QQ.1. Simultaneous QQ activities against AHL and AI-2 were exclusively demonstrated for protein 91_5/E6_ORF1. BLAST analysis revealed that this ORF showed homology to an integrase of *Pseudoalteromonas spp*. (74 % identity on nucleotide level, 98 % identity on protein level) (Fig. 3). Although QS interference has been demonstrated and verified, the underlying QQ remains highly speculative and has to be elucidated by further analysis.

**Fig. 3:**
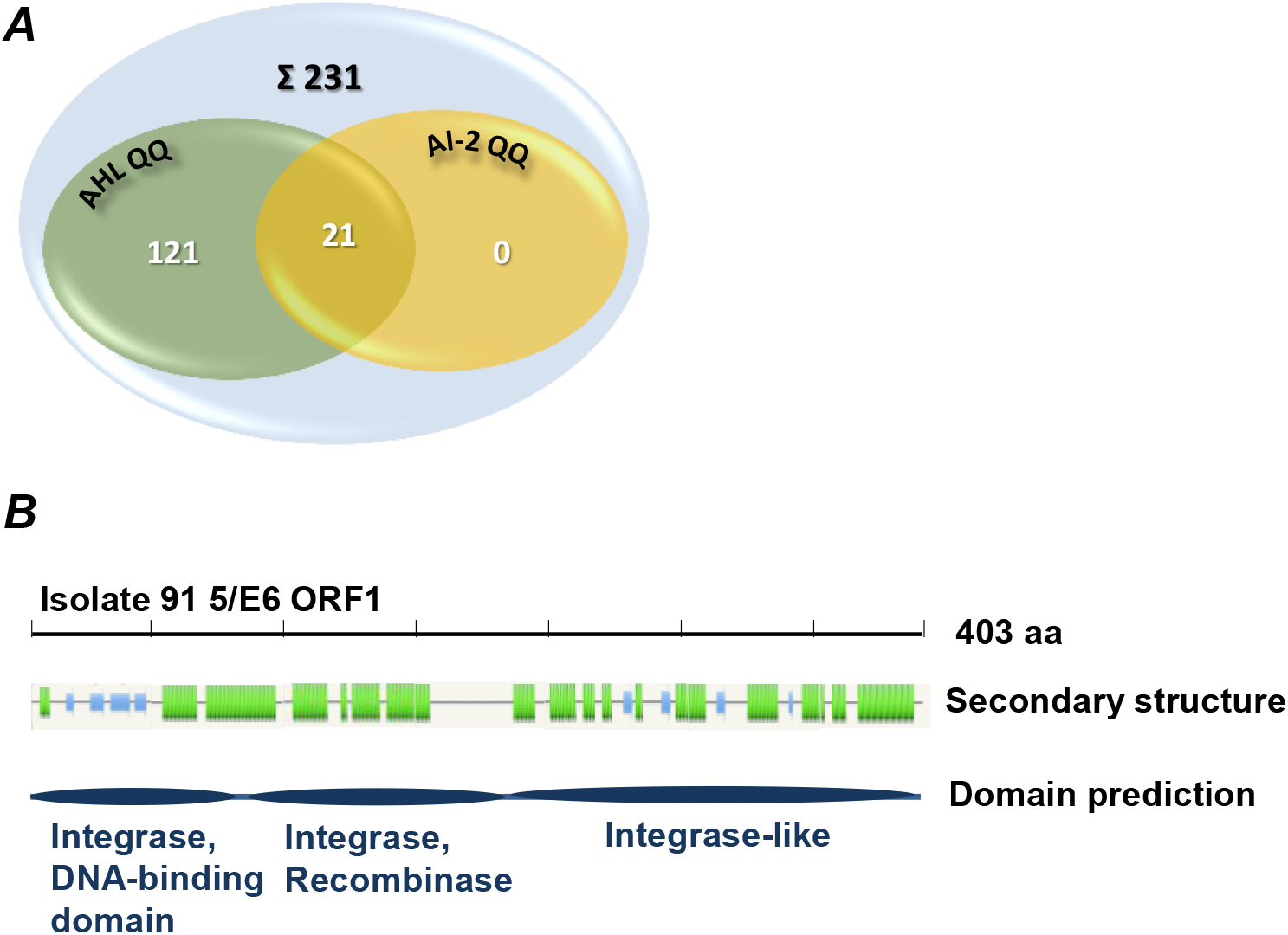
Quorum quenching activities of isolates. (***A***) Venn diagram illustrates QQ activities of isolates against acyl-homoserine lactones (AHL) and autoinducer-2 (AI-2). (***B***) QQ proteins were analyzed using secondary structure prediction software Phyre2 (http://www.sbg.bio.ic.ac.uk/phyre2/) with following visual output blue, strand; green, helix. InterProScan (http://www.ebi.ac.uk/Tools/pfa/iprscan/) was used for functional protein analysis.

In summary, with over 50 % we identified a high frequency of QS-interfering bacteria from the marine environment, which mainly interfere AHL communication primarily present in the marine environment (59, 60). These findings are in in accordance with recent reports on the occurrence of marine bacteria with AHL-QQ activities in pelagic and marine surface-associated communities (61, 62). In the study of Romero *et al*. (61), they demonstrated that 15 % of cultivable bacterial strains isolated from different marine microbial communities interfered AHL communication, a significantly higher percentage than that reported for soil isolates (2-5 %) (63, 64). QS and QQ seems to have an important ecological role in marine environments, particularly in dense microbial communities on biological surfaces. Many marine bacteria have evolved efficient strategies to combat neighboring bacteria, ultimately affecting the microbial community dynamics and patterns. Moreover, the identified high frequencies of QQ activities in marine bacteria disclose the enormous potential of marine bacteria as rich source for the identification of novel, biotechnologically relevant compounds, in particular QQ compounds for the inhibition of harmful biofilms (26, 60, 65).

## Acknowledgements

We thank Charlotte Eich, Johannes Effe, and Tim Quäschling for their experimental support in isolating and characterizing the bacteria. We thank the Institute of Clinical Molecular Biology in Kiel, Germany, for providing Sanger sequencing, supported in part by the DFG Cluster of Excellence Inflammation at Interfaces and Future Ocean. The work has received financial support from the DFG as part of the CRC1182 "Origin and function of metaorganisms”.

## Supplement

**Table S1: Primers used for QQ-ORF amplification and sequence analysis**. Underlined parts are added restriction sites for directed cloning.

**Tab. S2: Bacteria isolated from *Aurelia aurita, Mnemiopsis leidyi* and ambient seawater**. Bacteria were isolated by classical enrichment on agar plates and taxonomically classified based on full length 16S rRNA gene sequences. QQ activities of isolates are stated in (−) no activity, (+) low, (++) mid, and (+++) high activity against acyl-homoserine lactone (AHL) and autoinducer-2 (AI-2).

### Supplemental dataset

Sequence data of identified putative QQ-ORFs of fosmid clone 5/E6 originating from the genomic library of isolate 91 (*Pseudoalteromonas issachenkonii*)

