## Supplemental tables 1 and 2 for "Evaluating the quorum quenching potential of bacteria associated to *Aurelia aurita* and *Mnemiopsis leidyi*"

**Tab. S1: Primers used for QQ-ORF amplification and sequence analysis.** Underlined parts are added restriction sites for directed cloning.

| Primer | Sequence 5´→ 3´ |
|---|---|
| 91_5/E6_ORF1_for | GAATTCATGAACTTTACAGACAAATCAATTAAGGC |
| 91_5/E6_ORF1_rev | TCTAGATTACAATACTAAGCTTACTTTATGGC |
| 91_5/E6_ORF2_for | GAATTCATGCGTGGTACCCTAAACATTGC |
| 91_5/E6_ORF2_rev | AAGCTTTTACAGACCTCCCTTGAGATACCGC |
| 91_5/E6_ORF3_for | GAATTCATGAGCAACAATAAAGACACAGTGC |
| 91_5/E6_ORF3_rev | AAGCTTTTAATGAGCAACAATAAAGACACAG |

Supplement

**Tab. S2: Bacteria isolated from *Aurelia aurita*, *Mnemiopsis leidyi* and ambient seawater.** Bacteria were isolated by classical enrichment on agar plates and taxonomically classified based on full length 16S rRNA gene sequences. QQ activities of isolates are stated in (-) no activity, (+) low, (++) mid, and (+++) high activity against acyl-homoserine lactone (AHL) and autoinducer-2 (AI-2).

| **Isolate** | **origin** | **best homologue based on 16S full length rRNA gene (Accession No., identity)** | **taxonomic classification** | | | | **colony morphology** | **QQ activity** | |
|---|---|---|---|---|---|---|---|---|---|
| | | | **phylum** | **class** | **order** | **family** | | **AHL** | **AI-2** |
| **1** | *A. aurita* medusa Baltic Sea | *Pseudomonas stutzeri* (JX177727.1, 99%) | Proteobacteria | Gammaproteobacteria | Pseudomonadales | Pseudomonadaceae | yellowish-white, smeary | - | - |
| **2** | *A. aurita* medusa Baltic Sea | *Pseudomonas stutzeri* (JN228326.1, 99%) | Proteobacteria | Gammaproteobacteria | Pseudomonadales | Pseudomonadaceae | orange, round | ++ | - |
| **3** | *A. aurita* medusa Baltic Sea | *Pseudomonas segetis* (AY770691.2, 99%) | Proteobacteria | Gammaproteobacteria | Pseudomonadales | Pseudomonadaceae | white, round, smeary | - | - |
| **4** | *A. aurita* medusa Baltic Sea | *Pseudoalteromonas sp.* 191Z-13 (JX310231.1, 99%) | Proteobacteria | Gammaproteobacteria | Pseudomonadales | Pseudomonadaceae | white, smeary | +++ | - |
| **5** | *A. aurita* medusa Baltic Sea | Uncultured bacterium clone ncd1928c11c1 (JF165581.1, 93%) | n.a. | n.a. | n.a. | n.a. | white with orange spots, smeary | +++ | - |
| **6** | *A. aurita* medusa Baltic Sea | *Vibrio alginolyticus* (KP698605.1, 99%) | Proteobacteria | Gammaproteobacteria | Vibrionales | Vibrionaceae | white with orange spots, smeary | - | - |
| **7** | *A. aurita* medusa Baltic Sea | *Vibrio sp. S54CA* (KF188533.1, 98%) | Proteobacteria | Gammaproteobacteria | Vibrionales | Vibrionaceae | white, smeary | - | - |
| **8** | *A. aurita* medusa Baltic Sea | *Pseudomonas stutzeri* (JN228326.1, 99%) | Proteobacteria | Gammaproteobacteria | Pseudomonadales | Pseudomonadaceae | yellow, round | ++ | - |
| **9** | *A. aurita* medusa Baltic Sea | *Pseudomonas stutzeri* (JX177727.1, 99%) | Proteobacteria | Gammaproteobacteria | Pseudomonadales | Pseudomonadaceae | yellow, roundish | + | - |
| **10** | *A. aurita* medusa Baltic Sea | *Pseudomonas sp.* BJGMM-B54 (JQ716254.1, 99%) | Proteobacteria | Gammaproteobacteria | Pseudomonadales | Pseudomonadaceae | yellow, roundish | ++ | - |
| **11** | *A. aurita* medusa Baltic Sea | *Pseudomonas stutzeri* (JX177727.1, 99%) | Proteobacteria | Gammaproteobacteria | Pseudomonadales | Pseudomonadaceae | yellow, roundish | ++ | - |
| **12** | *A. aurita* medusa Baltic Sea | *Alteromonas genoviensis* (FJ040187.1, 99%) | Proteobacteria | Gammaproteobacteria | Alteromonadales | Alteromonadaceae | light yellow, round | ++ | - |

Supplement

| | | | | | | | | | |
|---|---|---|---|---|---|---|---|---|---|
| **13** | *A. aurita* medusa Baltic Sea | *Bacillus subtilis* (JX188065.1, 98%) | Firmicutes | Bacilli | Bacillales | Bacillaceae | white, oval | - | - |
| **14** | *A. aurita* medusa Baltic Sea | *Microbacterium lacticum* strain 2833 (EU714346.1, 99%) | Actinobacteria | Actinobacteria | Actinomycetales | Microbacteriaceae | white, smeary | - | - |
| **15** | *A. aurita* medusa Baltic Sea | *Sulfitobacter sp*. QD214-NF102 (KC689801.1, 99%) | Proteobacteria | Alphaproteobacteria | Rhodobacterales | Rhodobacteraceae | light orange in centre, round | +++ | + |
| **16** | *A. aurita* medusa Baltic Sea | *Micrococcus sp*. 3455 (KP345948.1, 99%) | Actinobacteria | Actinobacteria | Actinomycetales | Micrococcaceae | white, round | - | - |
| **17** | *A. aurita* medusa Baltic Sea | *Bacillus cereus* (KF624695.1, 98%) | Firmicutes | Bacilli | Bacillales | Bacillaceae | white, roundish | - | - |
| **18** | *A. aurita* medusa Baltic Sea husbandry | *Staphylococcus aureus* (CP011528.1, 99%) | Firmicutes | Bacilli | Bacillales | Staphylococcaceae | yellow, roundish-smeary | ++ | + |
| **19** | *A. aurita* medusa Baltic Sea husbandry | *Bacillus cereus* (DQ339684.1, 99%) | Firmicutes | Bacilli | Bacillales | Bacillaceae | white, smeary | - | - |
| **20** | *A. aurita* medusa Baltic Sea husbandry | *Phaeobacter gallaeciensis* (NR_027609.1, 82%) | Proteobacteria | Alphaproteobacteria | Rhodobacterales | Rhodobacteraceae | white, smeary | + | - |
| **21** | *A. aurita* medusa Baltic Sea husbandry | *Cobetia amphilecti* (NR_113404.1, 99%) | Proteobacteria | Gammaproteobacteria | Halomonadaceae | Cobetia | white-orange, smeary | + | - |
| **22** | *A. aurita* medusa Baltic Sea husbandry | *Pseudolateromonas sp*. MACL07 (EF198247.1, 99%) | Bacteroidetes | Flavobacteriia | Flavobacteriales | Flavobacteriaceae | orange, round | ++ | - |
| **23** | *A. aurita* medusa Baltic Sea husbandry | *Sulfitobacter sp*. S7-80 (KU999998.1, 99%) | Proteobacteria | Alphaproteobacteria | Rhodobacterales | Rhodobacteraceae | white, round | ++ | - |
| **24** | *A. aurita* medusa Baltic Sea husbandry | *Pseudoalteromonas sp*. MACL07 (EF198247.1, 99%) | Proteobacteria | Gammaproteobacteria | Pseudomonadales | Pseudomonadaceae | light orange, round | + | - |
| **25** | *A. aurita* medusa Baltic Sea husbandry | *Pseudoalteromonas issachenkonii* (JQ799065.1, 99%) | Proteobacteria | Gammaproteobacteria | Pseudomonadales | Pseudomonadaceae | white, round | ++ | - |
| **73** | *A. aurita* polyp | *Enterococcus casseliflavus* | Firmicutes | Bacilli | Lactobacillales | Enterococcaceae | white, roundish-smeary | + | - |

| | | | | | | | | | |
|---|---|---|---|---|---|---|---|---|---|
| | Baltic Sea husbandry | (KJ571214.1, 99%) | | | | | | | |
| **74** | *A. aurita* polyp Baltic Sea husbandry | *Micrococcus sp.* 3723 (KP345967.1, 99%) | Actinobacteria | Actinobacteria | Actinomycetales | Micrococcaceae | light yellow, round | - | - |
| **75** | *A. aurita* polyp Baltic Sea husbandry | *Arthrobacter davidanieli* (AF099202.1, 99%) | Actinobacteria | Actinobacteria | Actinomycetales | Micrococcaceae | white, round | - | - |
| **76** | *A. aurita* polyp Baltic Sea husbandry | *Bacillus mycoides* (CP009692.1, 98%) | Firmicutes | Bacilli | Bacillales | Bacillaceae | white, irregular | - | - |
| **77** | *A. aurita* polyp Baltic Sea husbandry | *Vibrio ordalii* (KC884626.1, 99%) | Proteobacteria | Gammaproteobacteria | Vibrionales | Vibrionaceae | white-brownish, smeary | ++ | - |
| **78** | *A. aurita* polyp Baltic Sea husbandry | *Sulfitobacter sp*. DG885 (AY258079.1, 99%) | Proteobacteria | Alphaproteobacteria | Rhodobacterales | Rhodobacteraceae | white, smeary | ++ | - |
| **79** | *A. aurita* polyp Baltic Sea husbandry | *Olleya marilimosa* strain KMM6714 (KC247324.1, 99%) | Bacteroidetes | Flavobacteriia | Flavobacteriales | Flavobacteriaceae | orange, round | +++ | + |
| **80** | *A. aurita* polyp Baltic Sea husbandry | *Vibrio anguillarum* (JX966409.1, 99%) | Proteobacteria | Gammaproteobacteria | Vibrionales | Vibrionaceae | white-brown, smeary | +++ | - |
| **81** | *A. aurita* polyp Baltic Sea husbandry | *Vibrio anguillarum* (JX966409.1, 99%) | Proteobacteria | Gammaproteobacteria | Vibrionales | Vibrionaceae | yellow, round | +++ | ++ |
| **82** | *A. aurita* polyp Baltic Sea husbandry | *Arthrobacter sp*. MB182 (JF706644.1, 99%) | Actinobacteria | Actinobacteria | Actinomycetales | Micrococcaceae | white, round | + | - |
| **83** | *A. aurita* polyp Baltic Sea husbandry | *Gordonia terrae* (KM113032.1, 97%) | Actinobacteria | Actinobacteria | Actinomycetales | Nocardiaceae | light orange, round | - | - |
| **84** | *A. aurita* polyp Baltic Sea husbandry | *Arthrobacter sp*. MB182 (JF706644.1, 100%) | Actinobacteria | Actinobacteria | Actinomycetales | Micrococcaceae | white, round (tiny) | ++ | - |
| **85** | *A. aurita* polyp Baltic Sea husbandry | *Staphylococcus warneri* (LC035464.1, 99%) | Firmicutes | Bacilli | Bacillales | Staphylococcaceae | white-yellow, round | + | - |

| | | | | | | | | | |
|---|---|---|---|---|---|---|---|---|---|
| **86** | *A. aurita* polyp Baltic Sea husbandry | Alpha proteobacterium C45 (AB302365.1, 96%) | Proteobacteria | Alphaproteobacteria | n.a. | n.a. | white, smeary | - | - |
| **87** | *A. aurita* polyp Baltic Sea husbandry | *Staphylococcus sp. C34* (JX482523.1, 99%) | Firmicutes | Bacilli | Bacillales | Staphylococcaceae | brownish, round | + | + |
| **88** | *A. aurita* polyp Baltic Sea husbandry | *Paracoccus sp*. UBF-P7 (JX239761.1, 99%) | Proteobacteria | Alphaproteobacteria | Rhodobacterales | Rhodobacteraceae | white, smeary | - | - |
| **89** | *A. aurita* polyp Baltic Sea husbandry | Alpha proteobacterium C45 (AB302365.1, 99%) | Proteobacteria | Alphaproteobacteria | n.a. | n.a. | white, round | - | - |
| **90** | *A. aurita* polyp Baltic Sea husbandry | *Pseudomonas putida* (FJ577648.1, 96%) | Proteobacteria | Gammaproteobacteria | Pseudomonadales | Pseudomonadaceae | white-yellow, round | ++ | ++ |
| **91** | *A. aurita* polyp Baltic Sea husbandry | *Pseudoalteromonas issachenkonii* (JQ799065.1, 98%) | Proteobacteria | Gammaproteobacteria | Pseudomonadales | Pseudomonadaceae | white-brownish, smeary | +++ | + |
| **92** | *A. aurita* polyp Baltic Sea husbandry | *Pseudomonas putida* (GU191929.1, 99%) | Proteobacteria | Gammaproteobacteria | Pseudomonadales | Pseudomonadaceae | white, round | ++ | - |
| **93** | *A. aurita* polyp Baltic Sea husbandry | *Pseudomonas sp. GA87* (AB934380.1, 99%) | Proteobacteria | Gammaproteobacteria | Pseudomonadales | Pseudomonadaceae | white, round with dent | ++ | - |
| **94** | *A. aurita* polyp Baltic Sea husbandry | *Pseudomonas monteilii* (KP056325.1, 99%) | Proteobacteria | Gammaproteobacteria | Pseudomonadales | Pseudomonadaceae | orange, roundish | ++ | - |
| **95** | *A. aurita* polyp North Sea husbandry | *Streptococcus infantis* (GU561389.1, 98%) | Firmicutes | Bacilli | Lactobacillales | Streptococcaceae | white, round | - | - |
| **96** | *A. aurita* polyp North Sea husbandry | *Enterococcus casseliflavus* (KJ571214.1, 99%) | Firmicutes | Bacilli | Lactobacillales | Enterococcaceae | white, round | + | - |
| **97** | *A. aurita* polyp North Sea husbandry | *Shewanella sp*. W3-18-1 (CP000503.1, 99%) | Proteobacteria | Betaproteobacteria | Alteromonadales | Oxalobacteraceae | yellowish, smeary | + | + |

Supplement

| | | | | | | | | | |
|---|---|---|---|---|---|---|---|---|---|
| **98** | *A. aurita* polyp North Sea husbandry | *Olleya marilimosa* strain KMM6714 (KC247324.1, 99%) | Bacteroidetes | Flavobacteriia | Flavobacteriales | Flavobacteriaceae | yellow/orange, smeary | - | - |
| **99** | *A. aurita* polyp North Sea husbandry | *Pseudoalteromonas issachenkonii* (JQ799065.1, 99%) | Proteobacteria | Gammaproteobacteria | Pseudomonadales | Pseudomonadaceae | light orange, round | ++ | + |
| **100** | *A. aurita* polyp North Sea husbandry | *Sulfitobacter sp*. DG885 (AY258079.1, 99%) | Proteobacteria | Alphaproteobacteria | Rhodobacterales | Rhodobacteraceae | white-yellow, smeary | +++ | - |
| **101** | *A. aurita* polyp North Sea husbandry | *Pseudoalteromonas issachenkonii* (JQ799065.1, 99%) | Proteobacteria | Gammaproteobacteria | Pseudomonadales | Pseudomonadaceae | very light orange, smeary | - | - |
| **102** | *A. aurita* polyp North Sea husbandry | *Alteromonas sp*. SN2 (KJ781946.1, 99%) | Proteobacteria | Gammaproteobacteria | Alteromonadales | Alteromonadaceae | white, brownish in center, round-smeary | + | - |
| **103** | *A. aurita* polyp North Sea husbandry | *Pseudoalteromonas ruthenica* (NR_025140.1, 98-99%) | Proteobacteria | Gammaproteobacteria | Pseudomonadales | Pseudomonadaceae | brownish, smeary | +++ | - |
| **104** | *A. aurita* polyp North Sea husbandry | *Ruegeria mobilis* (HQ338148.1, 99%) | Proteobacteria | Alphaproteobacteria | Rhodobacterales | Rhodobacteraceae | light, orange, smeary | ++ | - |
| **105** | *A. aurita* polyp North Sea husbandry | *Shewanella basaltis* (KC534403.1, 99%) | Proteobacteria | Betaproteobacteria | Alteromonadales | Oxalobacteraceae | white-yellow, smeary | - | |
| **106** | *A. aurita* polyp North Sea husbandry | *Hymenobacter psychrophilus* (NR_117214.1, 98%) | Bacteroidetes | Cytophagia | Cytophagales | Hymenobacteraceae | orange-pink, round | - | - |
| **107** | *A. aurita* polyp North Sea husbandry | *Luteococcus japonicas* (NR_119351.1, 99%) | Actinobacteria | Actinobacteria | Actinomycetales | Propionibacteriaceae | orange, round, smeary | + | - |
| **108** | *A. aurita* polyp North Sea husbandry | *Chryseobacterium hominis* (JX100820.1, 98-99%) | Bacteroidetes | Flavobacteriia | Flavobacteriales | Flavobacteriaceae | white, round | + | - |
| **109** | *A. aurita* polyp North Sea husbandry | *Rhodococcus erythropolis* PR4 (CP011295.1, 100%) | Actinobacteria | Actinobacteria | Actinomycetales | Nocardiaceae | white, roundish-smeary | - | - |

| | | | | | | | | | |
|---|---|---|---|---|---|---|---|---|---|
| **110** | *A. aurita* polyp North Sea husbandry | *Enterococcus casseliflavus* (KJ571214.1, 98-99%) | Firmicutes | Bacilli | Lactobacillales | Enterococcaceae | translucent, smeary | - | - |
| **111** | *A. aurita* polyp North Sea husbandry | *Brevibacterium frigoritolerans* (JF411310.1, 99%) | Actinobacteria | Actinobacteria | Actinomycetales | Brevibacteriaceae | white, round | +++ | - |
| **112** | *A. aurita* polyp North Sea husbandry | *Rhodococcus sp.* B126 (KJ781946.1, 99%) | Actinobacteria | Actinobacteria | Actinomycetales | Nocardiaceae | white, smeary | ++ | + |
| **113** | *A. aurita* polyp North Sea husbandry | *Sulfitobacter sp*. 132Z-6 (JX310150.1, 99%) | Proteobacteria | Alphaproteobacteria | Rhodobacterales | Rhodobacteraceae | white, smeary | - | - |
| **114** | *A. aurita* polyp North Atlantic husbandry | *Pseudomonas stutzeri* (HM137032.1, 99%) | Proteobacteria | Gammaproteobacteria | Pseudomonadales | Pseudomonadaceae | white-yellow, round | ++ | - |
| **115** | *A. aurita* polyp North Atlantic husbandry | *Moraxella osloensis* (AB643593.1, 99%) | Proteobacteria | Gammaproteobacteria | Halobacteriales | Moraxellaceae | white, round | - | - |
| **116** | *A. aurita* polyp North Atlantic husbandry | *Enterococcus casseliflavus* (KJ571214.1, 99%) | Firmicutes | Bacilli | Lactobacillales | Enterococcaceae | white, round | - | - |
| **117** | *A. aurita* polyp North Atlantic husbandry | *Ruegeria mobilis* (HQ338132.1, 99%) | Proteobacteria | Alphaproteobacteria | Rhodobacterales | Rhodobacteraceae | pink, smeary | +++ | - |
| **118** | *A. aurita* polyp North Atlantic husbandry | *Micrococcus sp*. 3723 (KP345967.1, 96%) | Actinobacteria | Actinobacteria | Actinomycetales | Micrococcaceae | white, round | - | - |
| **119** | *A. aurita* polyp North Atlantic husbandry | *Pseudoalteromonas issachenkonii* (JQ799065.1, 99%) | Proteobacteria | Gammaproteobacteria | Pseudomonadales | Pseudomonadaceae | light orange, round | ++ | - |
| **120** | *A. aurita* polyp North Atlantic husbandry | *Pseudoalteromonas sp*. MACL07 (EF198247.1, 99%) | Proteobacteria | Gammaproteobacteria | Pseudomonadales | Pseudomonadaceae | white-brownish, round | ++ | - |
| **121** | *A. aurita* polyp North Atlantic husbandry | *Olleya sp.* MOLA 14 (AM990790.1, 99%) | Bacteroidetes | Flavobacteriia | Flavobacteriales | Flavobacteriaceae | light orange, round | ++ | - |
| **122** | *A. aurita* polyp North | *Glaciecola sp.* KMM 6755 | Proteobacteria | Gammaproteobacteria | Alteromonadales | Alteromonadaceae | white, smeary | - | - |

| | | | | | | | | | |
|---|---|---|---|---|---|---|---|---|---|
| | Atlantic husbandry | (KF273912.1, 99%) | | | | | | | |
| **123** | *A. aurita* polyp North Atlantic husbandry | *Luteococcus japonicus* (NR_119351.1, 99%) | Actinobacteria | Actinobacteria | Actinomycetales | Propionibacteriaceae | white, round | + | - |
| **124** | *A. aurita* polyp North Atlantic husbandry | *Staphylococcus epidermidis* (FJ030635.1, 97%) | Firmicutes | Bacilli | Bacillales | Staphylococcaceae | white, smeary | - | - |
| **125** | *A. aurita* polyp North Atlantic husbandry | *Sulfitobacter sp*. DG885 (AY258079.1, 99%) | Proteobacteria | Alphaproteobacteria | Rhodobacterales | Rhodobacteraceae | white, round | - | - |
| **126** | *A. aurita* polyp North Atlantic husbandry | *Sulfitobacter sp*. QD214-NF102 (KC689801.1, 99%) | Proteobacteria | Alphaproteobacteria | Rhodobacterales | Rhodobacteraceae | white-orange, round | +++ | - |
| **127** | *A. aurita* polyp North Atlantic husbandry | *Staphylococcus succinus subsp. casei* (NR_037053.1, 99%) | Firmicutes | Bacilli | Bacillales | Staphylococcaceae | orange, roundish | +++ | - |
| **128** | *A. aurita* polyp North Atlantic husbandry | *Rhodococcus sp. B126* (KJ781946.1, 98%) | Actinobacteria | Actinobacteria | Actinomycetales | Nocardiaceae | white, orange, round-smeary | +++ | - |
| **129** | *A. aurita* polyp North Atlantic husbandry | *Staphylococcus aureus* (CP011528.1, 99%) | Firmicutes | Bacilli | Bacillales | Staphylococcaceae | white-yellow, roundish | - | - |
| **130** | *A. aurita* polyp North Atlantic husbandry | *Pseudomonas pachastrellae* (EU603457.1, 99%) | Proteobacteria | Gammaproteobacteria | Pseudomonadales | Pseudomonadaceae | white, roundish | + | - |
| **131** | *A. aurita* polyp North Atlantic husbandry | *Rhodococcus sp*. FXJ8.222 (KM507704.1, 99%) | Actinobacteria | Actinobacteria | Actinomycetales | Nocardiaceae | translucent, round | +++ | - |
| **51** | *M. leidyi* Baltic Sea | Uncultured bacterium clone nck64c11c1 (KF072560.1, 92%) | n.a. | n.a. | n.a. | n.a. | yellow, roundish, smeary | - | - |
| **52** | *M. leidyi* Baltic Sea | *Micrococcus endophyticus* (JQ659309.1, 99%) | Actinobacteria | Actinobacteria | Actinomycetales | Micrococcaceae | light yellow, round | - | - |
| **53** | *M. leidyi* Baltic Sea | *Alteromonas sp*. 2c3 (AJ294361.1, 99%) | Proteobacteria | Gammaproteobacteria | Alteromonadales | Alteromonadaceae | white, roundish-smeary | ++ | - |
| **54** | *M. leidyi* Baltic Sea | *Lacinutrix sp*. JR-M6 (KJ461692.1, 99%) | Bacteroidetes | Flavobacteriia | Flavobacteriales | Flavobacteriaceae | white-orange, round | ++ | - |

Supplement

| | | | | | | | | | |
|---|---|---|---|---|---|---|---|---|---|
| **55** | *M. leidyi* Baltic Sea | *Marinomonas hwangdonensis* (NR_109448.1, 98%) | Proteobacteria | Gammaproteobacteria | Oceanospirillales | Oceanospirillaceae | translucent, round | +++ | - |
| **56** | *M. leidyi* Baltic Sea | *Colwellia sp.*BSs20120 (EU330346.1, 99%) | Proteobacteria | Gammaproteobacteria | Alteromonadales | Colwelliaceae | light orange, round | - | - |
| **57** | *M. leidyi* Baltic Sea | *Olleya marilimosa* strain KMM6714 (KC247324.1, 99%) | Bacteroidetes | Flavobacteriia | Flavobacteriales | Flavobacteriaceae | translucent, round | +++ | + |
| **58** | *M. leidyi* Baltic Sea | *Rhodococcus sp.* ZS342 (JX428878.1, 99%) | Actinobacteria | Actinobacteria | Actinomycetales | Nocardiaceae | orange, roundish-smeary | - | - |
| **59** | *M. leidyi* Baltic Sea | *Microbacterium sp.* CDR2P2B2 (KJ567128.1, 99%) | Actinobacteria | Actinobacteria | Actinomycetales | Microbacteriaceae | white, round | - | - |
| **60** | *M. leidyi* Baltic Sea | *Alteromonas sp.* 2c3 (AJ294361.1, 99%) | Proteobacteria | Gammaproteobacteria | Alteromonadales | Alteromonadaceae | white, round, smeary | ++ | - |
| **61** | *M. leidyi* Baltic Sea | *Phaeocystidibacter luteus* (HQ434766.1, 99%) | Bacteroidetes | Flavobacteriia | Flavobacteriales | Cryomorphaceae | dark orange, oval | - | - |
| **62** | *M. leidyi* Baltic Sea | *Sagittula sp.* BG-9-E2 (KF560336.1, 89%) | Proteobacteria | Alphaproteobacteria | Rhodobacterales | Rhodobacteraceae | yellowish, oval | - | - |
| **63** | *M. leidyi* Baltic Sea | *Microbacterium oxydans* (KP136285.1, 99%) | Actinobacteria | Actinobacteria | Actinomycetales | Microbacteriaceae | white, round | - | - |
| **213** | *M. leidyi* Baltic Sea | *Vibrio sp.* (MF975618.1, 98%) | Proteobacteria | Gammaproteobacteria | Vibrionales | Vibrionaceae | yellowish, round | +++ | + |
| **215** | *M. leidyi* Baltic Sea | *Pseudomonas sp.* (KM461109.1, 99%) | Proteobacteria | Gammaproteobacteria | Pseudomonadales | Pseudomonadaceae | white-yellow, round | + | - |
| **218** | *M. leidyi* Baltic Sea | *Pseudoclavibacter sp.* (KY074321.1, 97%) | Actinobacteria | Actinobacteria | Actinomycetales | Microbacteriaceae | yellow, round | - | - |
| **219** | *M. leidyi* Baltic Sea | *Pseudoalteromonas tunicata* (KY319053.1, 99%) | Proteobacteria | Gammaproteobacteria | Pseudomonadales | Pseudomonadaceae | violet, round | +++ | - |
| **221** | *M. leidyi* Baltic Sea | *Aeromonas salmonicida* (HG941669.1, 98%) | Proteobacteria | Gammaproteobacteria | Aeromonadales | Aeromonadaceae | translucent, round | ++ | - |
| **222** | *M. leidyi* Baltic Sea | *Marinomonas pontica* (NR_042965.1, 100%) | Proteobacteria | Gammaproteobacteria | Oceanospirillales | Oceanospirillaceae | translucent, round | +++ | - |
| **223** | *M. leidyi* Baltic Sea | Uncultured *Alteromonas sp.* clone G9UC_PoM_0m_07 (KP076503.1, 98%) | Proteobacteria | Gammaproteobacteria | Alteromonadales | Alteromonadaceae | white with black center, round | - | - |
| **224** | *M. leidyi* Baltic Sea | *Pseudoalteromonas sp.* (KY671155.1, 97%) | Proteobacteria | Gammaproteobacteria | Pseudomonadales | Pseudomonadaceae | white, round | +++ | + |
| **225** | *M. leidyi* Baltic Sea | Acinetobacter sp. (JX266367.1, 99%) | Proteobacteria | Gammaproteobacteria | Halobacteriales | Moraxellaceae | white, spreading | - | - |

Supplement

| | | | | | | | | | |
|---|---|---|---|---|---|---|---|---|---|
| **228** | *M. leidyi* Baltic Sea | *Shewanella sp.* (KX230028.1, 96%) | Proteobacteria | Betaproteobacteria | Alteromonadales | Oxalobacteraceae | yellow, round | ++ | ++ |
| **229** | *M. leidyi* Baltic Sea | *Shewanella sp.* (JQ867500.1, 99%) | Proteobacteria | Betaproteobacteria | Alteromonadales | Oxalobacteraceae | red, round | +++ | + |
| **232** | *M. leidyi* Baltic Sea | *Pseudoalteromonas sp.* (JQ406678.1, 99%) | Proteobacteria | Gammaproteobacteria | Pseudomonadales | Pseudomonadaceae | orange, round | + | - |
| **233** | *M. leidyi* Baltic Sea | *Shewanella sp.* (KX531009.1, 99%) | Proteobacteria | Betaproteobacteria | Alteromonadales | Oxalobacteraceae | white, round | - | - |
| **234** | *M. leidyi* Baltic Sea | *Pseudoalteromonas sp.* (EU935585.1, 99%) | Proteobacteria | Gammaproteobacteria | Pseudomonadales | Pseudomonadaceae | black, round | +++ | - |
| **235** | *M. leidyi* Baltic Sea | *Shewanella sp.* (KX531009.1, 99%) | Proteobacteria | Betaproteobacteria | Alteromonadales | Oxalobacteraceae | yellow, round | ++ | - |
| **237** | *M. leidyi* Baltic Sea | Shewanella sp. (JQ867500.1, 99%) | Proteobacteria | Betaproteobacteria | Alteromonadales | Oxalobacteraceae | red, round | - | - |
| **240** | *M. leidyi* Baltic Sea | *Pseudoalteromonas sp.* (HQ882787.1, 99%) | Proteobacteria | Gammaproteobacteria | Pseudomonadales | Pseudomonadaceae | yellow, round | +++ | - |
| **241** | *M. leidyi* Baltic Sea | *Pseudoalteromonas sp.* (EU330361.1, 99%) | Proteobacteria | Gammaproteobacteria | Pseudomonadales | Pseudomonadaceae | white, round | - | - |
| **242** | *M. leidyi* Baltic Sea | Uncultured bacterium clone A3_91 (MF113934.1, 99%) | n.a. | n.a. | n.a. | n.a. | translucent, round | - | - |
| **243** | *M. leidyi* Baltic Sea | *Pseudoalteromonas tunicata* (KY319053.1, 99%) | Proteobacteria | Gammaproteobacteria | Pseudomonadales | Pseudomonadaceae | black, round | + | - |
| **246** | *M. leidyi* Baltic Sea | *Pseudomonas sp.* (HQ844525.1, 97%) | Proteobacteria | Gammaproteobacteria | Pseudomonadales | Pseudomonadaceae | white-yellow, spreading | +++ | - |
| **247** | *M. leidyi* Baltic Sea | *Shewanella sp.* (JF825437.1, 98%) | Proteobacteria | Betaproteobacteria | Alteromonadales | Oxalobacteraceae | translucent-yellow, round | +++ | - |
| **248** | *M. leidyi* Baltic Sea | Uncultured bacterium clone Woods-Hole_a2237 (KF798527.1, 98%) | n.a. | n.a. | n.a. | n.a. | white-yellow, round | - | - |
| **249** | *M. leidyi* Baltic Sea | *Pseudoalteromonas sp.* (KT583320.1, 97%) | Proteobacteria | Gammaproteobacteria | Pseudomonadales | Pseudomonadaceae | black, round | ++ | - |
| **250** | *M. leidyi* Baltic Sea | *Pseudoalteromonas sp.* (FR821214.1, 99%) | Proteobacteria | Gammaproteobacteria | Pseudomonadales | Pseudomonadaceae | violet, round | +++ | - |
| **251** | *M. leidyi* Baltic Sea | *Pseudoalteromonas tunicata* (KY319053.1, 99%) | Proteobacteria | Gammaproteobacteria | Pseudomonadales | Pseudomonadaceae | white, round | - | - |
| **254** | *M. leidyi* Baltic Sea | *Shewanella sp.* (KC247331.1, 99%) | Proteobacteria | Betaproteobacteria | Alteromonadales | Oxalobacteraceae | red, round | +++ | - |
| **255** | *M. leidyi* Baltic Sea | *Pseudoclavibacter sp*. (KM199858.1, 97%) | Actinobacteria | Actinobacteria | Actinomycetales | Microbacteriaceae | light yellow, round | - | - |

Supplement

| | | | | | | | | | |
|---|---|---|---|---|---|---|---|---|---|
| **256** | *M. leidyi* Baltic Sea | *Pseudoalteromonas tunicata* (KY319053.1, 99%) | Proteobacteria | Gammaproteobacteria | Pseudomonadales | Pseudomonadaceae | yellow, round | + | - |
| **260** | *M. leidyi* Baltic Sea | *Shewanella sp.* (KX692892.1, 99%) | Proteobacteria | Betaproteobacteria | Alteromonadales | Oxalobacteraceae | white-red, round | +++ | + |
| **261** | *M. leidyi* Baltic Sea | *Psychrobacter cryohalolentis* (KY405931.1, 98%) | Proteobacteria | Gammaproteobacteria | Halobacteriales | Moraxellaceae | white, spreading | - | - |
| **262** | *M. leidyi* Baltic Sea | *Marinomonas pontica* (NR_042965.1, 98%) | Proteobacteria | Gammaproteobacteria | Oceanospirillales | Oceanospirillaceae | translucent, round | +++ | - |
| **264** | *M. leidyi* Baltic Sea | *Exiguobacterium acetylicum* (MG490164.1, 99%) | Firmicutes | Bacilli | Bacillales | Bacilli | orange, round | - | - |
| **265** | *M. leidyi* Baltic Sea | *Pseudomonas sp.* (KY907020.1, 98%) | Proteobacteria | Gammaproteobacteria | Pseudomonadales | Pseudomonadaceae | yellow, round | + | - |
| **269** | *M. leidyi* Baltic Sea | *Phaeobacter daeponensis* (NR_044026.1, 99%) | Proteobacteria | Alphaproteobacteria | Rhodobacterales | Rhodobacteraceae | translucent-yellow, oval | - | - |
| **270** | *M. leidyi* Baltic Sea | *Bacillus sp.* (KF746902.1, 97%) | Firmicutes | Bacilli | Bacillales | Bacillaceae | black, round | - | - |
| **65** | *M. leidyi* Baltic Sea husbandry | *Pseudoalteromonas atlantica* (KP645203.1, 99%) | Proteobacteria | Gammaproteobacteria | Pseudomonadales | Pseudomonadaceae | white, smeary | + | + |
| **66** | *M. leidyi* Baltic Sea husbandry | *Alteromonas genoviensis* (FJ040187.1, 99%) | Proteobacteria | Gammaproteobacteria | Alteromonadales | Alteromonadaceae | white, round | ++ | - |
| **67** | *M. leidyi* Baltic Sea husbandry | *Shewanella sp*. KMM 6721 (KC247331.1, 99%) | Proteobacteria | Betaproteobacteria | Alteromonadales | Oxalobacteraceae | translucent, round | +++ | ++ |
| **68** | *M. leidyi* Baltic Sea husbandry | *Vibrio sp*. VibC-Oc-066 (KF577069.1, 99%) | Proteobacteria | Gammaproteobacteria | Vibrionales | Vibrionaceae | light orange, round | +++ | + |
| **69** | *M. leidyi* Baltic Sea husbandry | *Rhodobacter sp*. W402 (KF268394.1, 99%) | Proteobacteria | Alphaproteobacteria | Rhodobacterales | Rhodobacteraceae | orange, round | - | - |
| **70** | *M. leidyi* Baltic Sea husbandry | *Vibrio sp*. VibC-Oc-066 (KF577069.1, 99%) | Proteobacteria | Gammaproteobacteria | Vibrionales | Vibrionaceae | white, orange in center, round | ++ | - |
| **71** | *M. leidyi* Baltic Sea husbandry | *Ochrobactrum sp*. P1(2013) (KF987808.1, 99%) | Proteobacteria | Alphaproteobacteria | Rhizobiales | Brucellaceae | white, round | - | - |
| **72** | *M. leidyi* Baltic Sea husbandry | *Microbacterium sp.* MN2-1 (JQ396523.1, 99%) | Actinobacteria | Actinobacteria | Actinomycetales | Microbacteriaceae | yellowish-white, round | - | - |
| **217** | *M. leidyi* Baltic Sea husbandry | *Hydrogenophaga taeniospiralis* (AB795550.1, 99%) | Proteobacteria | Betaproteobacteria | Burkholderiales | Comamonadaceae | translucent-red, round | - | - |

Supplement

| | | | | | | | | | |
|---|---|---|---|---|---|---|---|---|---|
| **226** | *M. leidyi* Baltic Sea husbandry | *Pseudomonas sp.* (LC272923.1, 99%) | Proteobacteria | Gammaproteobacteria | Pseudomonadales | Pseudomonadaceae | light yellow, round | ++ | - |
| **227** | *M. leidyi* Baltic Sea husbandry | *Staphylococcus sp.* (MG162674.1, 99%) | Firmicutes | Bacilli | Bacillales | Staphylococcaceae | yellow-red, round | - | - |
| **239** | *M. leidyi* Baltic Sea husbandry | *Pseudoalteromonas sp.* (JX310130.1, 99%) | Proteobacteria | Gammaproteobacteria | Pseudomonadales | Pseudomonadaceae | white-yellow, round | + | - |
| **244** | *M. leidyi* Baltic Sea husbandry | *Thalassomonas sp.* (KC247368.1, 97%) | Proteobacteria | Gammaproteobacteria | Alteromonadales | Colwelliaceae | translucent-yellow, round | - | - |
| **257** | *M. leidyi* Baltic Sea husbandry | *Chryseobacterium sp.* (HQ911369.1, 97%) | Bacteroidetes | Flavobacteriia | Flavobacteriales | Flavobacteriaceae | yellow-red, round | ++ | - |
| **267** | *M. leidyi* Baltic Sea husbandry | *Alteromonas sp.* (KX989422.1, 99%) | Proteobacteria | Gammaproteobacteria | Alteromonadales | Alteromonadaceae | white, spreading | + | - |
| **214** | Ambient water Baltic Sea | *Serratia plymuthica* (KR611045.1, 99%) | Proteobacteria | Gammaproteobacteria | Enterobacteriales | Yersiniaceae | red, round | +++ | - |
| **230** | Ambient water Baltic Sea | *Pseudoalteromonas sp.* (KM979153.1, 99%) | Proteobacteria | Gammaproteobacteria | Pseudomonadales | Pseudomonadaceae | white, round | +++ | - |
| **231** | Ambient water Baltic Sea | *Pseudomonas sp.* (JF766700.1, 99%) | Proteobacteria | Gammaproteobacteria | Pseudomonadales | Pseudomonadaceae | white, spreading | - | - |
| **253** | Ambient water Baltic Sea | *Fictibacillus sp.* (KX033807.1, 98%) | Firmicutes | Bacilli | Bacillales | Bacillaceae | translucent yellow, round | - | - |
| **259** | Ambient water Baltic Sea | *Pseudoalteromonas sp.* (KF188488.1, 99%) | Proteobacteria | Gammaproteobacteria | Pseudomonadales | Pseudomonadaceae | red, round | ++ | - |
| **266** | Ambient water Baltic Sea | *Pseudoalteromonas sp.* (FR821212.1, 97%) | Proteobacteria | Gammaproteobacteria | Pseudomonadales | Pseudomonadaceae | translucent yellow, round | + | - |
| **132** | Artificial Seawater 18 PSU | *Pseudomonas anguilliseptica* (JX177685.1, 98-99%) | Proteobacteria | Gammaproteobacteria | Pseudomonadales | Pseudomonadaceae | light orange, smeary | - | - |
| **133** | Artificial Seawater 18 PSU | *Pseudomonas sp.* MBEF06 (AB733556.1, 97%) | Proteobacteria | Gammaproteobacteria | Pseudomonadales | Pseudomonadaceae | light orange, round | - | - |
| **134** | Artificial Seawater 18 PSU | *Pseudomonas anguilliseptica* (JX177685.1, 99%) | Proteobacteria | Gammaproteobacteria | Pseudomonadales | Pseudomonadaceae | light yellow, round | +++ | - |
| **135** | Artificial Seawater 18 PSU | *Moraxella sp.* CHZYR52 (AB905490.1, 99%) | Proteobacteria | Gammaproteobacteria | Halobacteriales | Moraxellaceae | white, round | ++ | - |

Supplement

| | | | | | | | | | |
|---|---|---|---|---|---|---|---|---|---|
| **136** | Artificial Seawater 18 PSU | *Pseudomonas anguilliseptica* (JX177685.1, 99%) | Proteobacteria | Gammaproteobacteria | Pseudomonadales | Pseudomonadaceae | white-yellowish, round | - | - |
| **137** | Artificial Seawater 18 PSU | *Sulfitobacter sp*. DG885 (AY258079.1, 99%) | Proteobacteria | Alphaproteobacteria | Rhodobacterales | Rhodobacteraceae | white, smeary | + | + |
| **138** | Artificial Seawater 18 PSU | *Celeribacter baekdonensis* (NR_117908.1, 99%) | Proteobacteria | Alphaproteobacteria | Rhodobacterales | Rhodobacteraceae | white, smeary | - | - |
| **139** | Artificial Seawater 18 PSU | *Alteromonas sp*. JAM-GA15 (AB526338.1, 99%) | Proteobacteria | Gammaproteobacteria | Alteromonadales | Alteromonadaceae | orange, round | ++ | - |
| **140** | Artificial Seawater 18 PSU | *Alteromonas sp*. SN2 (CP002339.1, 99%) | Proteobacteria | Gammaproteobacteria | Alteromonadales | Alteromonadaceae | white, smeary | ++ | - |
| **141** | Artificial Seawater 18 PSU | *Pseudomonas cuatrocienegasensis* (JN644592.1, 98%) | Proteobacteria | Gammaproteobacteria | Pseudomonadales | Pseudomonadaceae | yellow, round | - | - |
| **142** | Artificial Seawater 18 PSU | *Pseudomonas sp*. MBEF06 (AB733556.1, 99%) | Proteobacteria | Gammaproteobacteria | Pseudomonadales | Pseudomonadaceae | orange, smeary | + | - |
| **143** | Artificial Seawater 18 PSU | *Rhodococcus yunnanensis* (JN638050.1, 99%) | Actinobacteria | Actinobacteria | Actinomycetales | Nocardiaceae | white, round | - | - |
| **145** | Artificial Seawater 18 PSU | *Staphylococcus warneri* (LC035464.1, 99%) | Firmicutes | Bacilli | Bacillales | Staphylococcaceae | white, round | ++ | - |
| **146** | Artificial Seawater 18 PSU | *Staphylococcus pasteuri* (KP267845.1, 99%) | Firmicutes | Bacilli | Bacillales | Staphylococcaceae | white, round | - | - |
| **147** | Artificial Seawater 18 PSU | *Leisingera sp*. MA2-16 (KJ889016.1, 98%) | Proteobacteria | alphaproteobacteria | rhodobacterales | rhodobacteraceae | white, round | - | - |
| **148** | Artificial Seawater 18 PSU | *Alteromonas sp*. SN2 (CP002339.1, 99%) | Proteobacteria | Gammaproteobacteria | Alteromonadales | Alteromonadaceae | white, smeary | + | - |
| **149** | Artificial Seawater 18 PSU | *Staphylococcus saprophyticus* (KM095954.1, 99%) | Firmicutes | Bacilli | Bacillales | Staphylococcaceae | white, smeary | - | - |
| **150** | Artificial Seawater 18 PSU | *Staphylococcus warneri* (LC035464.1, 99%) | Firmicutes | Bacilli | Bacillales | Staphylococcaceae | white, round | - | - |
| **151** | Artificial Seawater 18 PSU | *Staphylococcus warneri* (LC035464.1, 99%) | Firmicutes | Bacilli | Bacillales | Staphylococcaceae | white line, round | - | - |
| **152** | Artificial Seawater 18 PSU | *Staphylococcus sp*. DVRSG-2 (KF779128.1, 99%) | Firmicutes | Bacilli | Bacillales | Staphylococcaceae | white line, smeary | - | - |

| | | | | | | | | | |
|---|---|---|---|---|---|---|---|---|---|
| **153** | Artificial Seawater 18 PSU | *Glaciecola sp*.DHVB6 (FJ848889.1, 99%) | Proteobacteria | Gammaproteobacteria | Alteromonadales | Alteromonadaceae | white, round | - | - |
| **154** | Artificial Seawater 18 PSU | *Alteromonas sp*. EM12a (HG004180.1, 99%) | Proteobacteria | Gammaproteobacteria | Alteromonadales | Alteromonadaceae | white, smeary | ++ | - |
| **155** | Artificial Seawater 18 PSU | *Marinobacter sp*. NP-1383C-30R (KJ914666.1, 99%) | Proteobacteria | Gammaproteobacteria | Alteromonadales | Alteromonadaceae | white-yellowish, round | +++ | - |
| **156** | Artificial Seawater 18 PSU | *Micrococcus sp*. 3723 (KP345967.1, 99%) | Actinobacteria | Actinobacteria | Actinomycetales | Micrococcaceae | yellow, round | - | - |
| **157** | Artificial Seawater 18 PSU | *Corynebacterium sp*. NML96-0244 (GU238410.1, 99%) | Actinobacteria | Actinomycetales | Corynebacteriaceae | Corynebacterium | white, round | - | - |
| **158** | Artificial Seawater 18 PSU | *Pseudomonas sp.* MT-1 (AP014655.1, 99%) | Proteobacteria | Gammaproteobacteria | Pseudomonadales | Pseudomonadaceae | light yellow, smeary | + | - |
| **159** | Artificial Seawater 18 PSU | *Pseudomonas sp*. MBEF06 (AB733556.1, 98-99%) | Proteobacteria | Gammaproteobacteria | Pseudomonadales | Pseudomonadaceae | light yellow, smeary | + | - |
| **160** | Artificial Seawater 18 PSU | *Staphylococcus aureus* (CP011528.1, 99%) | Firmicutes | Bacilli | Bacillales | Staphylococcaceae | yellow, round | - | - |
| **161** | Artificial Seawater 18 PSU | *Pseudomonas sp*. MBEF06 (AB733556.1, 98-99%) | Proteobacteria | Gammaproteobacteria | Pseudomonadales | Pseudomonadaceae | yellow-orange, smeary | - | - |
| **162** | Artificial Seawater 18 PSU | *Staphylococcus aureus* (CP011528.1, 99%) | Firmicutes | Bacilli | Bacillales | Staphylococcaceae | orange, smeary | - | - |
| **163** | Artificial Seawater 18 PSU | *Alteromonas sp*. JAM-GA15 (AB526338.1, 99%) | Proteobacteria | Gammaproteobacteria | Alteromonadales | Alteromonadaceae | orange, round | +++ | - |
| **164** | Artificial Seawater 18 PSU | *Celeribacter sp*. CY411 (KP201135.1, 99%) | Proteobacteria | Alphaproteobacteria | Rhodobacterales | Rhodobacteraceae | white, round | - | - |
| **165** | Artificial Seawater 18 PSU | *Celeribacter baekdonensis* (NR_117908.1, 99%) | Proteobacteria | Alphaproteobacteria | Rhodobacterales | Rhodobacteraceae | white with light yellow center, smeary | - | - |
| **166** | Artificial Seawater 18 PSU | *Celeribacter baekdonensis* (NR_117908.1, 99%) | Proteobacteria | Alphaproteobacteria | Rhodobacterales | Rhodobacteraceae | white, smeary | - | - |
| **167** | Artificial Seawater 18 PSU | *Pseudoalteromonas sp*. BSi20316 (DQ492738.1, 99%) | Proteobacteria | Gammaproteobacteria | Pseudomonadales | Pseudomonadaceae | white-yellowish, round | ++ | - |
| **168** | Artificial Seawater 18 PSU | *Salinibacterium sp.* ZS4-2 (FJ196007.1, 99%) | Actinobacteria | Actinobacteria | Actinomycetales | Microbacteriaceae | yellow, round | - | - |

Supplement

| | | | | | | | | | |
|---|---|---|---|---|---|---|---|---|---|
| **169** | Artificial Seawater 18 PSU | *Rhodococcus fascians* (FJ999590.1, 99%) | Actinobacteria | Actinobacteria | Actinomycetales | Nocardiaceae | orange, roundish | - | - |
| **170** | Artificial Seawater 18 PSU | *Pseudomonas stutzeri* strain QT 34 (HQ848122.1, 99%) | Proteobacteria | Gammaproteobacteria | Pseudomonadales | Pseudomonadaceae | yellow, round | +++ | - |
| **171** | Artificial Seawater 18 PSU | *Pseudomonas sp*. MBEF06 (AB733556.1, 98-99%) | Proteobacteria | Gammaproteobacteria | Pseudomonadales | Pseudomonadaceae | yellowish, round | ++ | - |
| **172** | Artificial Seawater 30 PSU | *Bacillus sp.* O-NR1 (JN613469.1, 99%) | Firmicutes | Bacilli | Bacillales | Bacillaceae | white, round | - | - |
| **173** | Artificial Seawater 30 PSU | *Salinibacterium amurskyense* strain y358 (KF306352.1, 99%) | Actinobacteria | Actinobacteria | Actinomycetales | Microbacteriaceae | light orange-white, round | - | - |
| **174** | Artificial Seawater 30 PSU | *Staphylococcus aureus subsp. aureus* SA268 (CP006630.1, 93%) | Firmicutes | Bacilli | Bacillales | Staphylococcaceae | white, round | - | - |
| **175** | Artificial Seawater 30 PSU | *Micrococcus sp.* 3723 (KP345967.1, 99%) | Actinobacteria | Actinobacteria | Actinomycetales | Micrococcaceae | orange, roundish-smeary | - | - |
| **176** | Artificial Seawater 30 PSU | *Halomonas boliviensis* (JX262399.1, 99%) | Proteobacteria | Gammaproteobacteria | Oceanospirillales | Halomonadaceae | light orange, round | - | - |
| **177** | Artificial Seawater 30 PSU | *Maribacter sp.* H24 (FJ903191.1, 98) | Bacteroidetes | Flavobacteriia | Flavobacteriales | Flavobacteriaceae | yellowish, smeary | + | - |
| **178** | Artificial Seawater 30 PSU | *Salinibacterium amurskyense* (KF306352.1, 98-99%) | Actinobacteria | Actinobacteria | Actinomycetales | Microbacteriaceae | yellowish, round | - | - |
| **179** | Artificial Seawater 30 PSU | *Bacillus sp.* Aza15 (JQ977243.1, 99%) | Firmicutes | Bacilli | Bacillales | Bacillaceae | white, round | - | - |
| **180** | Artificial Seawater 30 PSU | *Staphylococcus aureus* (CP011528.1, 99%) | Firmicutes | Bacilli | Bacillales | Staphylococcaceae | white, brownish center, round | +++ | - |
| **181** | Artificial Seawater 30 PSU | *Chryseobacterium sp.* WW-RP5 (KJ958497.1, 99%) | Bacteroidetes | Flavobacteriia | Flavobacteriales | Flavobacteriaceae | light orange, smeary | - | - |
| **182** | Artificial Seawater 30 PSU | *Microbacterium sp.* Cai-b5 (JX997907.1, 99%) | Actinobacteria | Actinobacteria | Actinomycetales | Microbacteriaceae | pink, round | - | - |
| **184** | Artificial Seawater 30 PSU | *Bacillus simplex* (KJ161409.1, 96%) | Firmicutes | Bacilli | Bacillales | Bacillaceae | white, brownish center, round | - | - |
| **185** | Artificial Seawater | *Alteromonas macleodii* (KP074899.1, 99%) | Proteobacteria | Gammaproteobacteria | Alteromonadales | Alteromonadaceae | orange, round | +++ | - |

| | | | | | | | | | |
|---|---|---|---|---|---|---|---|---|---|
| | 30 PSU | | | | | | | | |
| **186** | Artificial Seawater 30 PSU | *Alteromonas sp*. SCS1700m-1 (JX533655.1, 99%) | Proteobacteria | Gammaproteobacteria | Alteromonadales | Alteromonadaceae | brownish, round | +++ | - |
| **187** | Artificial Seawater 30 PSU | *Bacillus sp*. 7B-230 (KF441670.1, 99%) | Firmicutes | Bacilli | Bacillales | Bacillaceae | pink, smeary | - | - |
| **188** | Artificial Seawater 30 PSU | *Sulfitobacter pseudonitzschiae* strain H3 (KF006321.2, 99%) | Proteobacteria | Alphaproteobacteria | Rhodobacterales | Rhodobacteraceae | white, smeary | +++ | - |
| **189** | Artificial Seawater 30 PSU | *Bacillus vietnamensis* (KF933713.1, 100%) | Firmicutes | Bacilli | Bacillales | Bacillaceae | pink, round | - | - |
| **190** | Artificial Seawater 30 PSU | *Pseudoalteromonas sp.* 191Z-13 (JX310231.1, 99%) | Proteobacteria | Gammaproteobacteria | Pseudomonadales | Pseudomonadaceae | light-orange, round | - | - |
| **191** | Artificial Seawater 30 PSU | *Celeribacter sp*. CY411 (KP201135.1, 99%) | Proteobacteria | Alphaproteobacteria | Rhodobacterales | Rhodobacteraceae | white, round | - | - |
| **192** | Artificial Seawater 30 PSU | *Microbacterium sp*. JL1103 (DQ985063.1, 99%) | Actinobacteria | Actinobacteria | Actinomycetales | Microbacteriaceae | light orange, smeary | - | - |
| **193** | Artificial Seawater 30 PSU | *Brevibacterium frigoritolerans* (KJ767331.1, 99%) | Actinobacteria | Actinobacteria | Actinomycetales | Brevibacteriaceae | white, roundish | ++ | - |
| **194** | Artificial Seawater 30 PSU | *Cobetia amphilecti* (KP204120.1, 85%) | Proteobacteria | Gammaproteobacteria | Halomonadaceae | Cobetia | pink, round | + | - |
| **195** | Artificial Seawater 30 PSU | *Bacillus sp.* T1T (AM983464.1, 99%) | Firmicutes | Bacilli | Bacillales | Bacillaceae | translucent, round | - | - |
| **196** | Artificial Seawater 30 PSU | *Pseudomonas syringae pv. pisi* (KP211411.1, 99%) | Proteobacteria | Gammaproteobacteria | Pseudomonadales | Pseudomonadaceae | yellow, round | +++ | + |
| **197** | Artificial Seawater 30 PSU | *Pseudomonas pachastrellae* (KM460937.1, 99%) | Proteobacteria | Gammaproteobacteria | Pseudomonadales | Pseudomonadaceae | white, round | - | - |
| **199** | Artificial Seawater 30 PSU | *Maribacter sp*. H24 (FJ903191.1, 93) | Bacteroidetes | Flavobacteriia | Flavobacteriales | Flavobacteriaceae | white, round, smeary in high density | - | - |
| **200** | Artificial Seawater 30 PSU | *Pseudoalteromonas sp*. AB293f (FR821202.1, 99%) | Proteobacteria | Gammaproteobacteria | Pseudomonadales | Pseudomonadaceae | grey-white, round | ++ | - |
| **201** | Artificial Seawater 30 PSU | *Cobetia amphilecti* (NR_113404.1, 99%) | Proteobacteria | Gammaproteobacteria | Halomonadaceae | Cobetia | white, round | ++ | - |

Supplement

| | | | | | | | | | |
|---|---|---|---|---|---|---|---|---|---|
| **202** | Artificial Seawater 30 PSU | *Maribacter sp.* H24 (FJ903191.1, 99%) | Bacteroidetes | Flavobacteriia | Flavobacteriales | Flavobacteriaceae | white, round, small | - | - |
| **203** | Artificial Seawater 30 PSU | *Pseudoalteromonas espejiana* (KP204135.1, 99%) | Proteobacteria | Gammaproteobacteria | Pseudomonadales | Pseudomonadaceae | white, round, smeary in high density | +++ | - |
| **204** | Artificial Seawater 30 PSU | *Sulfitobacter sp*. QD214-NF102 (KC689801.1, 99%) | Proteobacteria | Alphaproteobacteria | Rhodobacterales | Rhodobacteraceae | white, round, smeary in high density | - | - |
| **205** | Artificial Seawater 30 PSU | *Pseudomonas sp*. MBTN3D1-a3 (JN975148.1, 89%) | Proteobacteria | Gammaproteobacteria | Pseudomonadales | Pseudomonadaceae | white, round | - | - |
| **206** | Artificial Seawater 30 PSU | *Bacillus mycoides* (CP009692.1, 99%) | Firmicutes | Bacilli | Bacillales | Bacillaceae | white-orange/brownish, smeary | - | - |
| **207** | Artificial Seawater 30 PSU | *Bacillus hwajinpoensis* (KJ009474.1, 99%) | Firmicutes | Bacilli | Bacillales | Bacillaceae | white-orange, smeary | - | - |
| **208** | Artificial Seawater 30 PSU | *Pseudoalteromonas issachenkonii* (JQ799065.1, 99%) | Proteobacteria | Gammaproteobacteria | Pseudomonadales | Pseudomonadaceae | white-yellowish, smeary | + | - |
| **209** | Artificial Seawater 30 PSU | *Brevibacterium frigoritolerans* (KF475857.1, 99%) | Actinobacteria | Actinobacteria | Actinomycetales | Brevibacteriaceae | white, roundish | - | - |
| **210** | Artificial Seawater 30 PSU | *Bacillus simplex* (KM817245.1, 99%) | Firmicutes | Bacilli | Bacillales | Bacillaceae | white, smeary | ++ | - |
| **212** | Artificial Seawater 30 PSU | *Fictibacillus phosphorivorans* (NR_118455.1, 99%) | Firmicutes | Bacilli | Bacillales | Bacillaceae | translucent, smeary | - | - |

Supplement

**Sequence data of identified putative QQ-ORFs of fosmid clone 5/E6 originating from the genomic library of isolate 91 (*Pseudoalteromonas issachenkonii*)**

I. 91_5/E6_ORF1

| Best homologue (based on aa sequence) | Protein | Accession No. | Identity |
|---|---|---|---|
| *Pseudoalteromonas sp*. TB41 | Integrase | WP_024602773.1 | 98 % |

>ORF1_nucleotide sequence

ATGAACTTTACAGACAAATCAATTAAGGCACTTAAGGCCAAAGAAAAACGTTACGTATTAACCGAGTCAGGTAACTAT
GGGGAAGGGCGTTTACAAATAAGGGTGAGTGAATCAGGCGCTAAAACGTTTCGGGTTCAGTATCACATAAACGGTAAG
CGCAAAGTAATTGGTCTTGGTAATTATCCTACCGTTGATCTTAAAAAAGCACGTAGTAAACATGCAAAAATAGCAGTC
TTATTAAGTGACAATATCGACCCGCAAGAGCATCAATTAGAAGCCCAAAAAATAGAGTTCGAATCATCAGCAAAGCGT
ACCATGTTACAAATGCTCGCTGACTTTAATGTATTCATAAGTACACGCTGGGCAGAGTCAACAATAGACCGAACTGAA
AAACTCATTAAAAGAAACATCACCCCGTTTATAAAACCCGAATTAATGCCCGACGAGTTCACCATAGATATGGCCCGC
GACATTATTTACCGTGTTTATAATCGTGGCGCAAAAGAACAAGCGCGACTAGTTCGCAGCATACTAATGAGCATATTA
AAATTTGCTATAGATTTTGATAACTCACCAGAGCAATACAAAAAGCCAAACCTATACGACATAAAAACAAATTTCATC
AGAGACATTAACTTTGAAACGCCAAAAAACAAAGGTGAACGCTGGTTAAGCGAGGCTGAACTAAAAAAAGTATGGAAT
GCAGACGACCTACCTTATTACACCCACCAATACATAAAACTGGCATTATTACTTGGTGGTCAGCGAGTAAATGAGGTT
TACGGCTCATACGTAAGTGACTTTGATTTAGAAAATAAAACTTTCACTATCCCCGCAAATCGTATCAAAGTACAACAA
CGGGGCGATCACATAGTGCCATTATGCGAAACCGCAATACCAATCATCCAAGAGCTAATTCAACAAGCAGGTAAAGCT
GGTCAAATGTTCCCGCATCGCGACAACCCAACAGCCACCGCCCATGTATCAACACTTCGAATGGCAATATTACGATGG
TGCGAAAAAAACAAAGTGCCAAACTTTAATCCCCGTGATCTACGCAGAACGTGTAAAACACTCATGGGCAAAGCAGGC
ATAGATAAAATAAACCGCGACATACTGCAGCAACACAACAAGTTTGATGTATCAAGTGTGCATTACGACAGATACGAC
TATATGAAAGAAAAACGCCAAAGCATTGAGGTGTGGGAAACTGCTTATGAATTGTGCCATAAAGTAAGCTTAGTATTG
TAA

>ORF1_amino acid sequence

MNFTDKSIKALKAKEKRYVLTESGNYGEGRLQIRVSESGAKTFRVQYHINGKRKVIGLGNYPTVDLKKARS
KHAKIAVLLSDNIDPQEHQLEAQKIEFESSAKRTMLQMLADFNVFISTRWAESTIDRTEKLIKRNITPFIK
PELMPDEFTIDMARDIIYRVYNRGAKEQARLVRSILMSILKFAIDFDNSPEQYKKPNLYDIKTNFIRDINF
ETPKNKGERWLSEAELKKVWNADDLPYYTHQYIKLALLLGGQRVNEVYGSYVSDFDLENKTFTIPANRIKV
QQRGDHIVPLCETAIPIIQELIQQAGKAGQMFPHRDNPTATAHVSTLRMAILRWCEKNKVPNFNPRDLRRT
CKTLMGKAGIDKINRDILQQHNKFDVSSVHYDRYDYMKEKRQSIEVWETAYELCHKVSLVL

II. 91_5/E6_ORF2

| Best homologue (based on aa sequence) | Protein | Accession No. | Identity |
|---|---|---|---|
| *Pseudoalteromonas* | Cation/H+ | WP_096038305.1 | 98 % |

| | |
|---|---|
| *teradonis* | antiporter |

>ORF2_nucleotide sequence

ATGCGTGGTACCGACACTTGGTTTTTATTAGTCGGTTTGACGGGTTTAACTACACTTTTATTTGGCGCTTACATCGCG
CTATTCAAACATGATTTAAAAGGCTTATTAGCCTATTCAACAATTAGTCATTTAGGCCTAATTACTCTATTACTCGGC
CTAGACACACAACTTGCAACCGTAGCCGCTATTTTTCATATTATTAACCATGCTACGTTTAAAGCGTCGTTATTTATG
GCCACGGGTATTATTGACCATGAAACCGGCACGCGTGATATGCGCAAACTCAATGGCATGTGGCGCTACTTACCTTAT
ACGGCCACATTAGCGATGGTGGCCGCTGCCGCTATGGCGGGTGTACCGCTATTAAATGGTTTCTTATCTAAAGAAATG
TTTTTTGCTGAAACACTGCATCAGCAAGTACTTGGCTCTATGTCGTGGTTAATTCCTGTGCTGGCAACCGTTGCAGGT
GCGCTGTCGGTAGCGTACTCATCTCGCTTTATTCATGACGTGTTCTTTAATGGTGAACCGATAGACTTACCACGCACC
CCTCATGAGGCGCCACGTTATATGCGTGTGCCTATCGAAATTTTAGTGGTGCTATGTATTTTAGTGGGTATTTTTCCG
CACTTTGCAGTAGATGGTATTTTATCGGCTGCCTCATTGGCCGTACTTGGCCAAGCTATGCCTGAGTACAAGCTAACT
ATTTGGCACGGTTTCAATTTACCTCTACTTATGAGTGGTATGGCGGTGATTGGTGGTTTATTTATTTACGTTAATCGT
AAGTACTTGTTCCAGTTTCAGGCGTCGTTACCTCCTTTTAACGCTAAAAAAATCTTCGAGCGTTTTTTAGCGGTTGTG
GTTAATTGGTGTCAAAACAAAATCCAATCAAACAGAAAATGGCTCATTACAACGCTATGTATTTATTATGTTAGGTGT
TGTATTGCTTGCCTCAGGTTGGCCATTATTTGA

>ORF2_amino acid sequence

MRGTDTWFLLVGLTGLTTLLFGAYIALFKHDLKGLLAYSTISHLGLITLLLGLDTQLATVAAIFHIINHATFKASLFM
ATGIIDHETGTRDMRKLNGMWRYLPYTATLAMVAAAAMAGVPLLNGFLSKEMFFAETLHQQVLGSMSWLIPVLATVAG
ALSVAYSSRFIHDVFFNGEPIDLPRTPHEAPRYMRVPIEILVVLCILVGIFPHFAVDGILSAASLAVLGQAMPEYKLT
IWHGFNLPLLMSGMAVIGGLFIYVNRKYLFQFQASLPPFNAKKIFERFLAVVVNWCQNKIQSNRKWLITTLCIYYVRC
CIACLRLAII

III. 91_5/E6_ORF3

| Best homologue (based on aa sequence) | Protein | Accession No. | Identity |
|---|---|---|---|
| *Pseudoalteromonas sp*. DSM 26666 | Cation/H+ antiporter | WP_09049398.1 | 100 % |

>ORF2_nucleotide sequence

ATGCACGGCGGCGGTAAACCAAAAACACGTGCAGCACTGCATTACGTTATTTTAAACTTGGTTGGCTCAAGTGTATTT
TTAATCGGCTTAGGTATTTTATATGGGGTATTGGGTACCCTAAACATTGCTGATATGGCTAATAAAGTACCGCAGTTA
ACTGGTGATGATGTTTATTTAGCTAAAGCGGGCGGGTTACTATTGCTTGTAGTATTTGCCCTTAAAAGTGCACTATTA
CCACTGCATTTATGGCTCCCAAATGCCTATTCAAGTGCGACACCTGTGGTTGCTGCATTATTCGCGATTATGACCAAA
GTAGGTGTTTACGCCACTTTACGCGTATATACAGTGGTATTTGGTGAGCAAGCCGGTGAACTTGAGCATATGGCGCAG
TCTTGGCTATGGGCGTTGGCCATTGCTACTATAGTGATTGGGGCAATCGGGGTGTTGGCAGCACAAGATTTACGTAAG
CTTACCGCAAACTTAGTACTGGTTTCTGTGGGTACTTTAGTGGCACTGGTTGCACTTCAAAATATAAACGCCACAGCT
GCACTATTGTATTATTTGGTGCACTCAACGTTAGTGACTGCAGCACTATTTTTACTGGCTGATTTAATTGCAACTCAG
CGAGGTAAAGCCGGCGACAGATTAGTTGGCGGACGCGCAGTTAAACAACCATTCTTGTTGGGAGCGTGTTTTATTATC
GCTGGACTAACCGTCATAGGTATGCCGCCACTTTCAGGTTTTGTAGGTAAAATTTGGATTTTAAAAACCACCTTAAAC
AGTGAGCAAGCACTGGTATTTTGGCCTGTTTACTTAATTATGAGCTTAGCGTTAATTGTTGCTATATCACGTGCAGGC
ACGAGTTTGTTTTGGGAGCATAAAGACAAAGGCGGTGAGGCTGGTGATGTTGCTAATGCACATCCGTTGCAAGTTATC

Supplement

GTACTTGTTGGCCTGCTCACAAGCTCAATACTGTTGGTTGTTTTTGGTGATTTGGCAACGCAATATGCACTTGAAACA
GCAACACAACTTCATGATATTAGTGGTGGTATTAATGCGGTACTCAAGGGAGGTCTGTAA

>ORF3_amino acid sequence

MHGGGKPKTRAALHYVILNLVGSSVFLIGLGILYGVLGTLNIADMANKVPQLTGDDVYLAKAGGLLLLVVFALKSALL
PLHLWLPNAYSSATPVVAALFAIMTKVGVYATLRVYTVVFGEQAGELEHMAQSWLWALAIATIVIGAIGVLAAQDLRK
LTANLVLVSVGTLVALVALQNINATAALLYYLVHSTLVTAALFLLADLIATQRGKAGDRLVGGRAVKQPFLLGACFII
AGLTVIGMPPLSGFVGKIWILKTTLNSEQALVFWPVYLIMSLALIVAISRAGTSLFWEHKDKGGEAGDVANAHPLQVI
VLVGLLTSSILLVVFGDLATQYALETATQLHDISGGINAVLKGGL
